# Seed microbiota legacy mitigates the effect of drought in wheat

**DOI:** 10.64898/2026.01.16.699829

**Authors:** Hamed Azarbad, Pranav M. Pande, Charlotte Giard-Laliberté, Asmaâ Agoussar, Jessica A. Dozois, Emmy L’Espérance, Luke D. Bainard, Julien Tremblay, Étienne Yergeau

## Abstract

Seeds carry epiphytes and endophytes microbial partners that can shape plant fitness. However, whether these microbial communities serve as transgenerational memory systems of parental stress remains largely unstudied. Here, we combined multi-year field rainfall manipulation experiments in eastern Canada with a greenhouse experiment in western Canada to test whether seed-associated microbiota transmit drought legacies across plant generations. In the field experiment, reduced rainfall initially decreased yield in the drought-sensitive wheat cultivar (AC Nass), but selected for distinct seed bacterial endophyte communities. In subsequent generations, plants whose seed microbiota retained compositional similarity to these drought-adapted communities showed enhanced yield stability under upcoming water stress. A transgenerational field test confirmed that daughter plants derived from drought-exposed parent plants maintained performance under water limitation, whereas those from wetter origins did not. In an independent greenhouse assay using seeds from Saskatchewan fields differing in long- and short-term irrigation history, AC Nass plants from water-stress legacy sites exhibited higher photosynthetic efficiency, water-use efficiency, and root bacterial diversity under drought. Together, these findings demonstrate that seed-associated microbiota act as ecological archives of stress history, transmitting drought legacies across generations in a cultivar and year-dependent manner.

## Introduction

Climate change is intensifying the frequency, duration, and severity of drought events worldwide (IPCC, 2023; Ohlert *et al*. 2025), placing unprecedented pressure on terrestrial plants to adapt across spatial and temporal scales. Plants function as the holobiont (Zilber-Rosenberg and Rosenberg 2008), relying on microbial partners that contribute to water-use regulation, stress signaling, and nutrient mobilization (Quiza, St-Arnaud and Yergeau 2015; Azarbad and Junker 2024; Compant *et al*. 2025). Increasing evidence shows that the plant-associated microbiota are not randomly assembled, but include vertically transmitted lineages inherited mainly through seeds (Bergna *et al*. 2018; Rodríguez *et al*. 2020; Abdelfattah *et al*. 2021, 2022; Becker and Cubeta 2024; Sulesky-Grieb *et al*. 2024). Whether drought can shape these seed-borne communities in a way that influences plant performance in subsequent generations remains poorly understood.

Among plant microbiota, those associated with seeds play an important role as a transmission vector, selectively retaining microbial taxa that may potentially influence early colonization, germination, and downstream interactions with soil and root communities (Berg and Raaijmakers 2018; Nelson 2018; Shahzad *et al*. 2018; War *et al*. 2023; Kumar *et al*. 2024). In the context of accelerating climate variability, vertically inherited seed microbiota may constitute an underappreciated mechanism through which parental exposure to drought or other abiotic stressors shapes the performance of next generations of plants. However, empirical evidence for such seed microbially mediated legacy effects remains scarce, particularly under different agricultural fields and across multiple plant generations. This is particularly important since plant microbiota can move along the soil–plant–human gut axis via food and modulate human gut community structure and immune-relevant metabolism, thus the seed microbiota has implications that extend from crop resilience to One Health (Berg *et al*. 2020; Ma, Cornadó and Raaijmakers 2025).

The legacy effect or stress memory of plant microbiota is supported across different ecosystems. Wheat plants developed greater root biomass when grown in soils with a history of water stress, indicating that microbial legacies in soil can buffer hosts against subsequent drought (Azarbad *et al*. 2018). Similarly, soil microbial communities exposed to chronic drought show enhanced functional resilience through stress-adapted gene enrichment (Broderick *et al*. 2025), while aquatic and synthetic communities recover more predictably after perturbation if pre-conditioned by earlier stress (Cairns *et al*. 2025; Xu *et al*. 2025). A similar mechanism could also be at play for seed-associated microbial communities, but has received less attention. Seeds carry endophytes, microbes residing within protected internal tissues, and epiphytes, microbes colonizing the exposed seed surface (Nelson 2018; Azarbad *et al*. 2022). These two compartments may differ in their environmental exposure, transmission pathways, and host filtering. However, it is not yet known whether such contrasting microbial compartments encode drought memory (or legacy), or whether they translate into measurable effects on the performance of the next generation of plants.

Here, we address these gaps using three experiments centered on drought-tolerant and sensitive wheat cultivars grown under field rainfall manipulations and annual reseeding. In the rainfall manipulation experiment (Experiment 1), a four-level rainfall gradient (100, 75, 50,25%) was applied over three seasons with annual reseeding, enabling separation of generational versus water-regime effects on seed-associated bacterial and fungal communities. In the transgenerational field test (Experiment 2), we asked whether the precipitation history of seed origin predicts performance when seeds are replanted under ambient versus sheltered (reduced rainfall) conditions in the subsequent year. Finally, in the irrigation history greenhouse experiment (Experiment 3), we evaluated the seed microbial legacies by comparing seeds produced under continuous vs. intermittent water stress history in a distinct agroecological region. For that, we studied plant physiological (assimilation and water-use efficiency) and morphological traits (biomass and height), as well as the rhizosphere microbiota under controlled greenhouse drought conditions. We hypothesized that drought leaves a microbial imprint in wheat seeds that is transmitted across generations, with compartment and microbial-specific patterns. Specifically, we expected that (i) seed epiphytes would exhibit stronger restructuring than endophytes under reduced rainfall, reflecting their greater exposure to environmental filtering; (ii) bacterial communities would respond more dynamically to drought history than fungi, and (iii) the strength of these seed microbiota legacies would depend on host cultivar, with the drought-sensitive bread wheat cultivar (*Triticum aestivum* cv. AC Nass) showing stronger response than the drought-tolerant durum wheat cultivar (*Triticum turgidum* subsp. durum cv. Strongfield). Finally, we hypothesized that these microbial legacies go beyond diversity or compositional shifts to produce measurable functional outcomes under subsequent stress, enhancing yield stability in the next plant generation derived from drought-history seeds.

## Results

### The rainfall manipulation experiment (Experiment 1)

#### (A) Precipitation, moisture dynamics, and yield responses

To determine how different precipitation treatments reshape wheat seed microbiota and influence plant performance across generations, we established a multi-year rainfall manipulation experiment in field plots in Laval, Québec (Fig. 1a). Beginning in 2016, two wheat species with contrasting drought sensitivities, *T. aestivum* cv. AC Nass (drought-sensitive) and *T. turgidum* subsp. durum cv. Strongfield (drought-tolerant), were grown under four precipitation regimes (100%, 75%, 50%, and 25% of ambient rainfall). At the end of each growing season (2016 and 2017), seeds were harvested from each plot and sown back into the same plots the following spring (Fig. 1b). Monthly precipitation varied across years, with 2017 exhibiting relatively higher rainfall during peak growing months (June–July), while, in general, 2018 experienced lower precipitation (Fig. 1c). Soil water content followed a clear treatment-dependent gradient, with drier treatments (25% and 50%) leading to significantly lower soil moisture levels in 2016 and 2017 than wetter treatments (75% and 100%; Fig. 1d). By 2018, soil moisture levels in all plots remarkedly declined compared to 2016 and 2017. Yield responses varied between the two wheat cultivars. In AC Nass, a significant effect was observed in 2016 (p = 0.039), where reduced rainfall negatively impacted yield, but this effect was no longer significant in 2017 (Fig. 1e). In contrast, for Strongfield yield remained unaffected by rainfall treatments across both years, supporting its drought tolerant characteristic (Fig. 1f).

**Figure 1.**
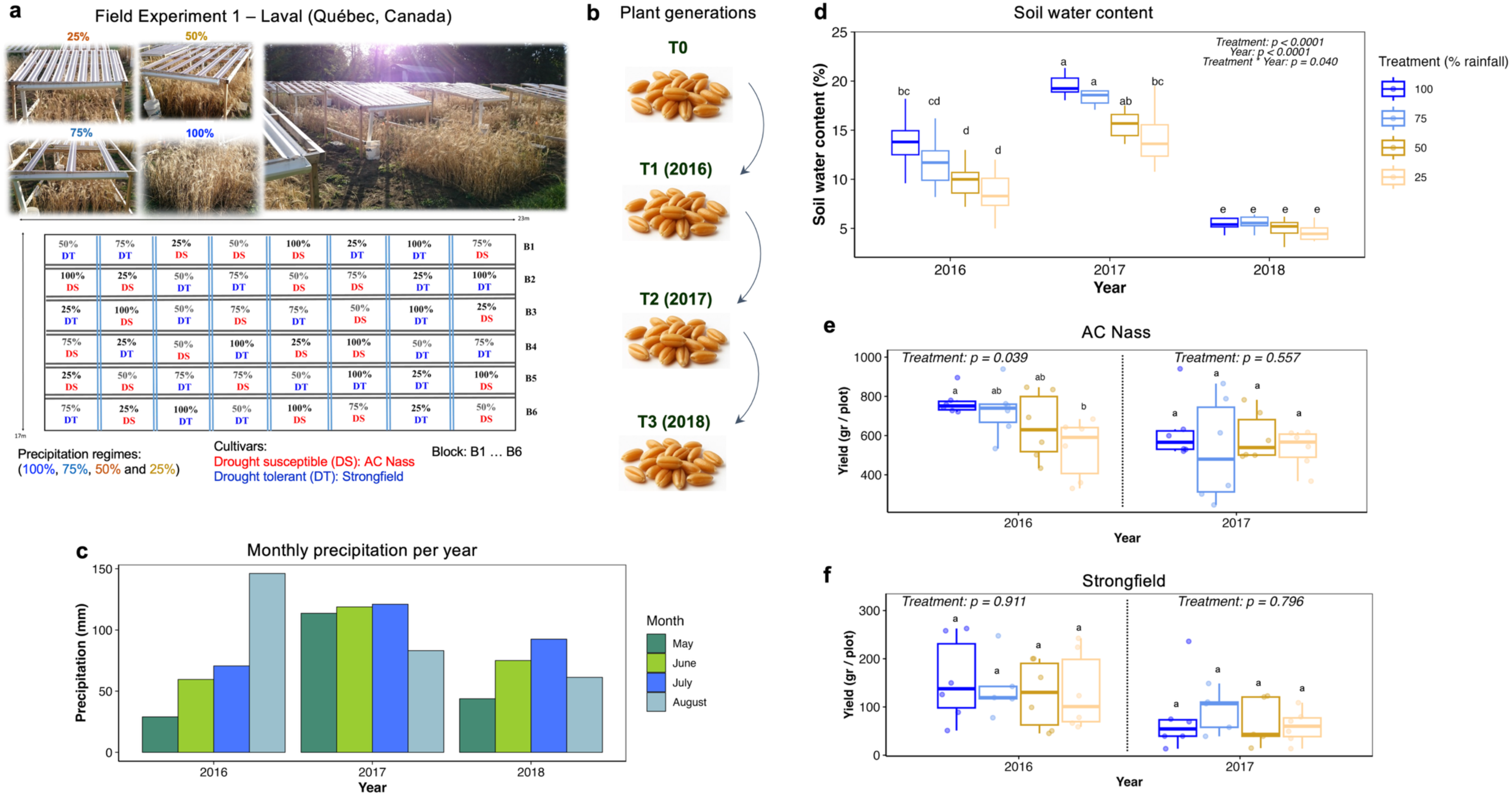
Rainfall manipulation experiment in Laval, Québec, and associated effects on soil water content and yield. **a**, Experimental design showing rainout shelters applying four rainfall regimes (100, 75, 50, 25% of ambient precipitation) in randomized plots with two cultivars (drought-susceptible AC Nass, drought-tolerant Strongfield). **b**, Generational design: pre-treatment seeds (T0) were sown in 2016 (T1), harvested, and re-sown in the same plots in 2017 (T2) and 2018 (T3). **c**, Monthly precipitation during the experimental period, with 2018 showing lower rainfall. **d**, Soil water content across treatments and years, showing significant effects of rainfall treatment, year, and their interaction. **e**, Grain yield of AC Nass, showing a significant treatment effect in 2016 but no effect in 2017. **f**, Grain yield of Strongfield, showing no significant treatment effect in either 2016 or 2017. Letters denote Tukey’s HSD groupings.

#### (B) Generational dynamics of bacterial and fungal seed microbiota under contrasting rainfall regimes

In AC Nass, generations (G) significantly influenced bacterial richness in the rhizosphere (χ² = 9.596, df = 2, p = 0.008), leaf (χ² = 32.642, df = 2, p < 0.0001), seed epiphytes (χ² = 18.363, df = 2, p < 0.0001), and seed endophytes (χ² = 13.968, df = 2, p = 0.001). Additionally, the interaction between rainfall and generations (T × G) significantly impacted bacterial richness in seed epiphytes (χ² = 31.160, df = 11, p = 0.001) and endophytes (χ² = 23.000, df = 11, p = 0.018). Strongfield showed no such responses (Supplementary Table S1). Given the stronger generational and precipitation-driven shifts in bacterial richness observed in AC Nass compared to Strongfield, we focused specifically on the seed-associated bacterial communities of AC Nass. In 2016 and 2017, the bacterial richness of seed epiphytes was relatively stable across rainfall treatments (Fig. 2a). In 2018, which was the driest year during the experimental periods, epiphytic bacterial richness declined sharply in the 25% treatment, while the 100% and 75% treatments retained greater diversity (Fig. 2a). In 2016, the richness of bacterial endophytes was higher in seeds from wet rainfall treatments (the 100% and 75%), compared with the dry treatments (50% and 25%: Fig. 2b; Fig. S1). Mean richness in dry treatments increased significantly (t = 3.70, p = 0.0007) from 2016 to 2017 by +13.0 ± 3.5, whereas richness in the wet treatments showed no change (+2.5 ± 3.6, p = 0.49; Fig. S1). By 2018, the lowest diversity was again observed in the 25% treatment, but still significantly higher than the first year of the experiment under the same treatment (Fig. 2b). Fungal seed epiphytes displayed moderate shifts across generations, and diversity reductions in the 25% treatment was only evident in 2018 (Fig. 2c), which was reflected in a significant treatment-by-generation interaction (χ² = 22.281, df = 11, p = 0.022; Supplementary Table S2). In contrast, seed endophytes were remarkably stable (Fig. 2d), and neither treatment nor its interaction with generation altered richness.

**Figure 2.**
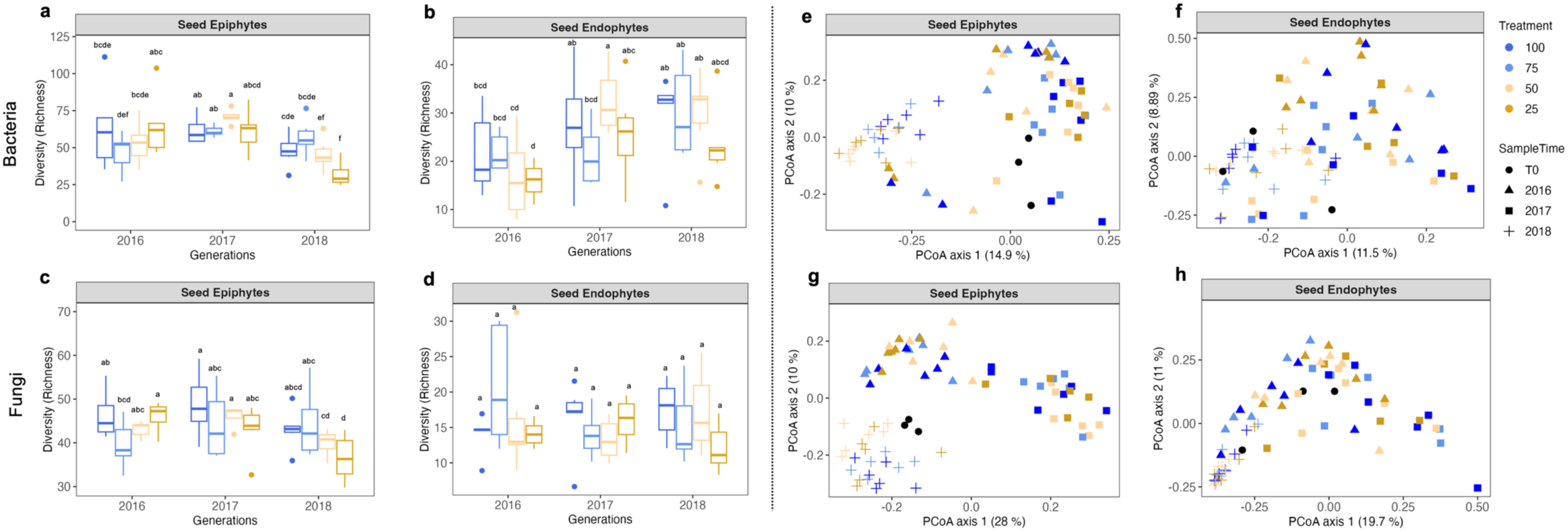
Multi-year dynamics of bacterial and fungal seed microbiota under rainfall manipulation in AC Nass. **a–b**, Bacterial richness of seed epiphytes (a) and seed endophytes (b) across rainfall treatments and generations (2016–2018). **c–d**, Fungal richness of seed epiphytes (c) and endophytes (d). **e–f**, Principal coordinate analysis (PCoA) of bacterial communities in seed epiphytes (e) and endophytes (f). **g–h**, PCoA of fungal communities in seed epiphytes (g) and endophytes (h).

### Microbial community structure

To assess the impact of rainfall manipulation and generational shifts on wheat-associated microbiota, we performed Permanova analyses separately for each cultivar (for bacteria: Supplementary Table S3; for fungi: Supplementary Table S4). Community-level differences were further visualized using PCoA (for bacteria, Fig. S2; for fungi, Fig. S3). Across compartments, generation (G) had the strongest effect on bacterial community structure for the rhizosphere, root, leaf, seed epiphytes, and endophytes (Supplementary Table S3). For seed bacterial epiphytes, in 2018, community divergence due to the treatment was most pronounced (Fig. 2e). As for seed bacterial endophytes, the interaction between rainfall treatment and generation (T × G) was significant (F = 1.358, p = 0.042; Supplementary Table S3; Fig. 2f). The rainfall treatment had a significant direct effect on shaping the community structure of fungal epiphytes (F = 1.806, p < 0.01) that was more apparent in 2018, where samples from drier conditions (25% and 50% rainfall) clustered separately from those under higher rainfall treatments (Fig. 2g). Fungal endophytes were shaped only by generation (Fig. 2h). For Strongfield, bacterial communities in the rhizosphere and seed epiphytes exhibited generational effects, whereas rainfall treatment had no significant impacts.

### Microbial community composition

We further examined changes in microbial community composition at the order level for all plant compartments in the case of both cultivars (bacterial: Fig. S4; fungal: Fig. S5). Given that seed-associated bacterial endophytes and fungal epiphytes of the drought-sensitive cultivar AC Nass exhibited the strongest responses to precipitation history (as shown by Permanova and PCoA analyses), we focused subsequent community-level analyses on ASVs level (bacterial: Fig. 3a; fungal: Fig. 3b) in these two compartments. ASV-level taxonomic profiling highlighted specific drought-responsive and generation-sensitive bacterial taxa (Fig. 3a). *Erwinia* (ASV_11) and *Escherichia–Shigella* (ASV_2) showed strong responses to both rainfall treatment and generation (p < 0.001), with *Erwinia* being enriched in 2018, particularly under drought conditions. In contrast, *Massilia* (ASV_13), although present in early years, declined under drought by 2018. *Pantoea* (ASV_31) displayed a distinct pattern where its relative abundance increased over time, emerging in 2018 across all rainfall treatments (Fig. 3a). As for fungi seed epiphytes, in 2016, *Alternaria* (ASV_1) accounted for the majority of the community across all rainfall regimes, with a lower relative abundance under dry conditions than under wet conditions (Fig. 3b). By 2017, fungal assemblages diversified, with reduced dominance of *Alternaria* and significant increases in *Rhodotorula* (ASV_11), *Moesziomyces* (ASV_12, 23), *Epicoccum* (ASV_24), and *Vishniacozyma* (ASV_25), yielding the most compositionally heterogeneous year. By 2018, communities contracted again: *Alternaria* (ASV_1) reasserted dominance in wetter treatments (100% and 75%), while drought-maintained seeds retained elevated *Cladosporium* (ASV_3), which showed a significant treatment × generation interaction (p < 0.05).

**Figure 3.**
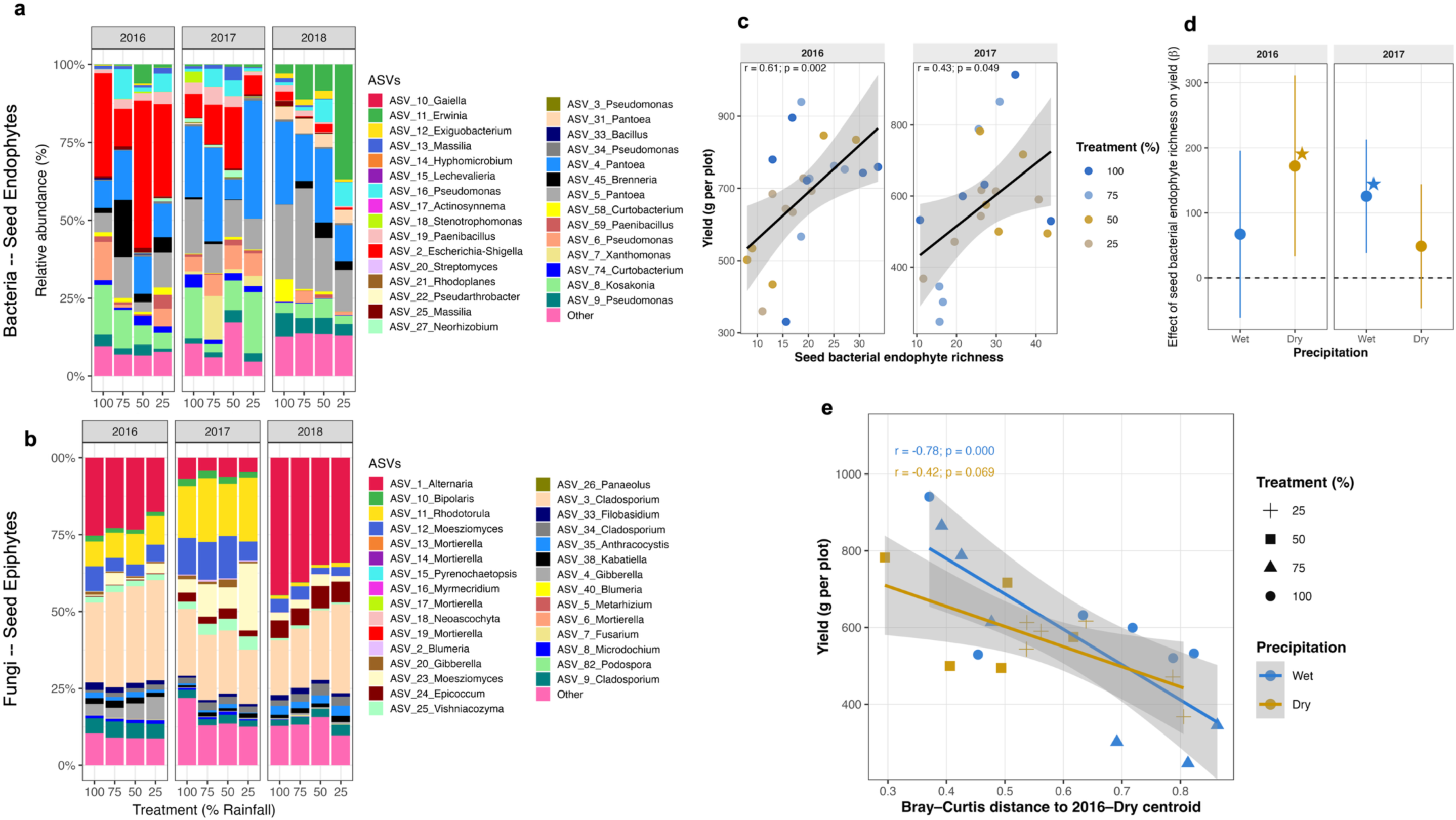
Seed endophyte richness and community composition predict yield resilience under drought history. **a**, Relative abundance of dominant bacterial ASVs associated with seed endophyte across rainfall treatments and years. **b**, Relative abundance of dominant fungal seed epiphyte ASVs. **c**, Relationship between seed bacterial endophyte richness and yield in 2016 and 2017. Each point represents an individual field plot, coloured by irrigation treatment (25–100%). Linear regressions (black lines ±95% confidence interval) illustrate the within-year associations between richness and yield. **d**, Effect of seed bacterial endophyte richness on yield derived from a linear mixed-effects model. The slopes represent the expected change in yield (grams per plot) for each one-standard-deviation increase in seed endophyte richness, after accounting for soil water content (SWC) and block effects. Points denote slope estimates for plants originating from wet (blue) and dry (gold) rainfall regimes in 2016, evaluated within each growing year (2016 and 2017). Stars indicate slopes significant after false-discovery-rate adjustment (q ≤ 0.05). The dashed horizontal line indicates a zero effect. **e**, Legacy effect of seed bacterial endophyte community composition on yield. Bray–Curtis distances were calculated between each 2017 sample and the centroid of the 2016–Dry community to quantify compositional similarity to the previous-year drought microbiota. Coloured regression lines show the relationship between compositional distance and yield for 2017 seed endophyte communities originating from wet (blue) or dry (gold) rainfall regimes, with shaded areas indicating 95% confidence intervals.

#### (C) The diversity and community composition of seed bacterial endophyte predict wheat yield under contrasting rainfall regimes

We next examined whether variations in seed-associated bacterial diversity and composition are related to changes in wheat yield under contrasting rainfall regimes for AC Nass. Based on the generational and treatment-dependent shifts described above, these analyses focused specifically on seed bacterial endophytes (for bacteria) and seed fungal epiphytes (for fungi), which displayed significant responses to rainfall treatment. As previously shown, seed bacterial endophyte richness increased significantly between 2016 and 2017 in plants from the drier rainfall treatments (Δ = +13.0; p < 0.001), whereas it remained unchanged under wetter treatments (Δ = +2.5; p = 0.49; Fig. S1). Interestingly, across both years, bacterial endophyte richness correlated positively with yield (2016: r = 0.61, p = 0.002; 2017: r = 0.43, p = 0.049; Fig. 3c). Linear mixed-effects models controlling for soil water content and block effects confirmed that the strength and direction of richness–yield relationships depended on rainfall regime (Fig. 3d). In 2016, richness had a positive effect on yield under dry treatment but not under wet conditions, whereas the pattern reversed in 2017, reflecting a year-dependent influence of diversity on plant performance. To assess whether the structure of the seed endophyte community itself carried a transgenerational “legacy” of drought adaptation, we quantified each 2017 community’s Bray–Curtis distance from the 2016-dry centroid (a compositional proxy for the drought-adapted microbiota). Yield declined with increasing distance from this dry reference (Fig. 3e), indicating that seed communities more compositionally similar to the prior-year drought assemblage conferred higher productivity. This relationship was particularly strong for plants originating from wet histories (β = –868.0; p < 0.001), but was not statistically significant for dry-origin plants (β = –474.6; p = 0.069). We did not detect any significant patterns between fungi data and the yield pattern.

### The transgenerational field test (Experiment 2): Precipitation history shapes yield stability under drought

The strong relationship between seed bacterial endophyte diversity, community composition, and yield observed in the Laval field experiment (Experiment 1) suggested that drought history may imprint a microbial legacy within seeds. Such legacy effects could, in principle, contribute to plant pre-adaptation in subsequent generations by transmitting microbiota better suited to specific moisture regimes. To test this hypothesis, we conducted a second field experiment using seeds harvested from the different rainfall treatments of Experiment 1 for a new generation of plants exposed to controlled drought and ambient conditions. For that, we grew wheat seeds collected from the 2016 and 2017 rainfall treatments under two water regimes: with shelter (reduced precipitation, ∼25 % of ambient) or without shelter (ambient rainfall; Fig. 4a). In 2017, no significant treatment effects were observed for either cultivar, suggesting that the immediate precipitation history of seeds did not influence yield when grown under varying water availability (Fig. 4b). However, the yield results of AC Nass in 2018, the driest year in the study period, revealed that seed-origin effects. Seeds from the wetter plot origin (100% and 75% precipitation in 2017) exhibited a significant yield reduction when grown under drought (with shelter) compared to ambient conditions (without shelter) (p < 0.05). In contrast, for seeds originating from dry-rainfall backgrounds (50% and 25% precipitation in 2017), no significant differences were detected between reduced and ambient precipitation (Fig. 4b). In contrast, Strongfield exhibited no significant differences across treatments, indicating its capacity for drought tolerance, independent of the contribution of its microbiota.

**Figure 4.**
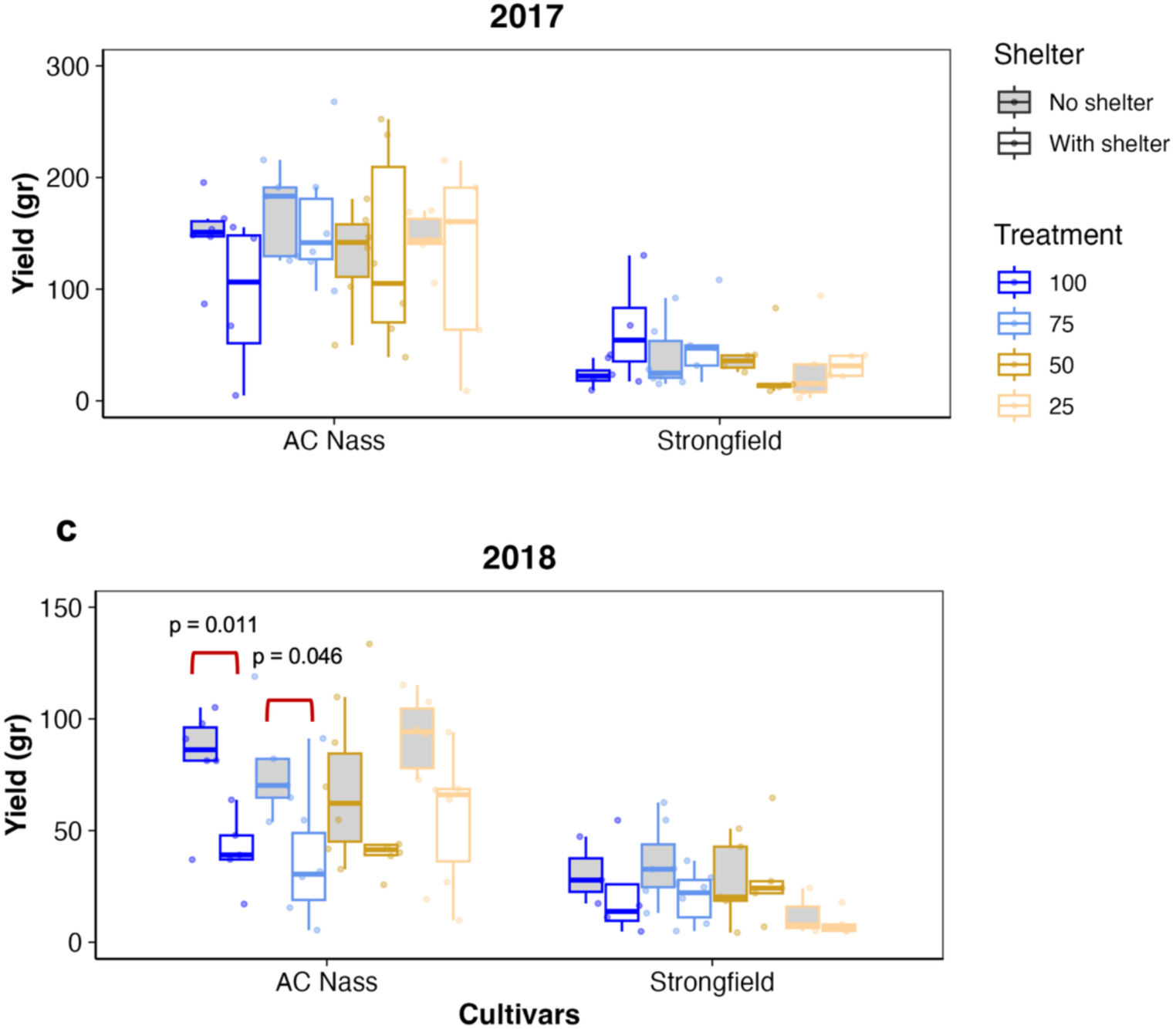
Transgenerational field experiment at Laval and effects of seed microbial precipitation history on yield under drought. Grain yield in 2017, showing no significant effect of shelter across precipitation histories for either cultivar. **c**, Grain yield in 2018, showing that AC Nass plants originating from wetter seed histories (100%, 75%) exhibited significant yield reductions under shelter compared with unsheltered controls, whereas seeds from drought histories (50%, 25%) showed no yield reductions.

### The irrigation history greenhouse experiment (Experiment 3): The legacy of field water stress history influences plant performance, rhizosphere bacterial diversity, and community composition under greenhouse drought

To assess whether seed legacy from contrasting field water stress histories influenced plant performance and associated microbiota, we conducted a greenhouse experiment using seeds harvested from a field trial in Saskatchewan (Fig. 5a). Plants from two wheat cultivars (AC Nass and Strongfield), originating from irrigated or non-irrigated field sites with continuous or intermittent water stress history (WSH), were grown under controlled soil water holding capacities (SWHC) of 50% or 20% (Fig. 5b).

**Figure 5.**
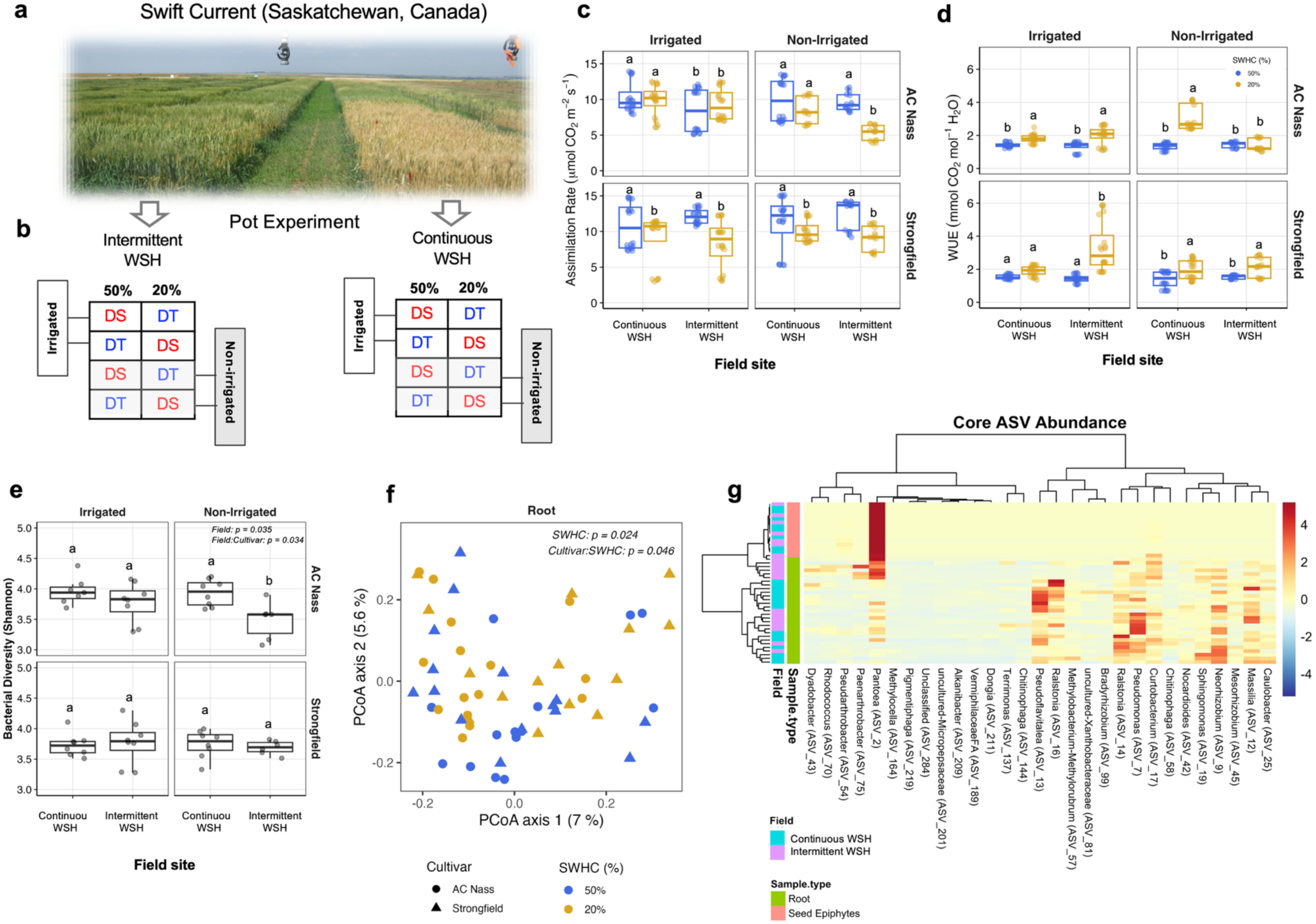
Irrigation history greenhouse experiment for validation of seed legacy effects from Saskatchewan field sites under controlled drought. **a**, Field plots in Swift Current contrasting irrigated and non-irrigated sites used as seed sources, with intermittent vs. continuous water stress history (WSH). **b**, Experimental design of the greenhouse validation, with seeds from irrigated and non-irrigated sites (intermittent or continuous WSH) grown under two soil water-holding capacities (SWHC; 50%, 20%) in AC Nass (drought-susceptible, DS) and Strongfield (drought-tolerant, DT). **c**, Net assimilation rates under greenhouse drought. **d**, Water-use efficiency (WUE) patterns. **e**, Shannon diversity of root bacterial communities, higher in AC Nass from continuous WSH sites compared with intermittent WSH controls. **f**, PCoA of root bacterial communities showing separation by SWHC and a significant cultivar × SWHC interaction. **g**, Heatmap of 29 core ASVs shared between seed epiphytes and root bacteria, showing compartmental and seed field source structuring.

#### Plant physiological and morphological responses

Across treatments, plants subjected to 20% SWHC exhibited significantly reduced assimilation rates (F = 38.95, P < 0.001) and increased water use efficiency (WUE; F = 150.5, p < 0.001), which are in line with expected reductions in photosynthetic carbon fixation and improved carbon gain per unit water loss under drought (Fig. 5c–d; Supplementary Table S5). Genotypic effect was also significant (assimilation: F = 23.08, p < 0.001; WUE: F = 9.00, p = 0.003), with AC Nass displaying greater sensitivity to field legacy than Strongfield. In AC Nass, assimilation was lowest in plants originating from non-irrigated sites with intermittent WSH, while seeds from non-irrigated sites with continuous WSH maintained higher assimilation under the same drought conditions (Fig. 5c). In contrast, Strongfield maintained relatively stable assimilation across field histories. As for WUE, AC Nass from continuous WSH fields under non-irrigated sites showed a significant increase in WUE under 20% SWHC, whereas plants from sites with intermittent WSH exhibited no significant changes than 50% SWHC (Fig. 5d). In contrast, Strongfield exhibited the opposite trend, with respect to legacy, in such a way that WUE increased under 20% SWHC were evident in plants from irrigated sites with intermittent WSH, whereas seeds from continuous WSH sites did not differ significantly between 20% and 50% SWHC (Supplementary Table S5; Fig. 5d). In terms of morphological responses, shoot biomass and plant height were significantly reduced under 20% SWHC (Shoot biomass: F = 56.24, p < 0.001; Plant height: F = 83.07, p < 0.001) and varied by cultivar (Shoot biomass: F = 36.73, p < 0.001; Plant height: F = 75.97, p < 0.001), but showed no significant main or interactive effects of field water stress history (Supplementary Table S5).

#### Root bacterial diversity and community composition

Root bacterial Shannon diversity was shaped by field history (p = 0.035) and its interaction with cultivar (p = 0.034; Fig. 5e). In AC Nass, plants from non-irrigated fields with intermittent WSH harbored the lowest root bacterial diversity, whereas those from continuous WSH fields maintained higher diversity. This pattern was not evident in Strongfield. PCoA of root bacterial communities revealed a significant effect of SWHC (*p* = 0.024) and a cultivar × SWHC interaction (*p* = 0.046) (Fig. 5f). This structuring was more pronounced in AC Nass, where 20% SWHC samples formed distinct clusters from 50%, compared to Strongfield. We further examined 29 core ASVs that were detected in both seed epiphytes and root microbiota (Fig. 5g). Clustering by sample type indicated a strong compartmental effect, with roots and seed epiphytes hosting distinct subsets of the core taxa. Within this compartmental separation, several ASVs varied in relative abundance according to field history. For example, *Pantoea* (ASV_2) tended to be more abundant in seed epiphytes, whereas *Bradyrhizobium* (ASV_14), *Pseudoflavitalea* (ASV_13), *Ralstonia* (ASV_13), and *Sphingomonas* (ASV_9) were more represented in roots from continuous WSH fields, and *Pseudomonas* (ASV_7) was more abundant in roots from intermittent WSH fields. No significant effect of experimental treatment was observed for the root fungi diversity and community patterns.

## Discussion

Across multiple experiments under different agroecological contexts, we showed that drought history leaves an ecological imprint on the wheat seed microbiota, which in turn modulates plant performance under subsequent water limitation. In the rainfall manipulation experiment, drought reduced yield in 2016 but simultaneously selected for distinct bacterial endophyte communities that became predictive of plant productivity in the AC Nass cultivar in the subsequent year. The following generation exhibited enhanced yield stability when its seed microbiota retained compositional similarity to the drought-adapted community of the previous season. The transgenerational field test confirmed that this microbial imprint can act as functional pre-adaptation in such a way that seeds originating from drought-exposed parental environments sustained yields under upcoming water limitation, while those from wet origins experienced yield declines. The irrigation history greenhouse experiment extended the results to a broader temporal scale where seeds produced in fields differing in decades-long irrigation history, yet themselves grown for only one season under common conditions, retained the imprint of that legacy. Plants derived from continuous WSH fields showed higher drought resilience compared to those from intermittent WSH fields. Importantly, the irrigation history experiment strongly suggests that the observed transgenerational effects are due to microorganisms and not to plant epigenetic mechanisms, as only the microorganisms could have transmitted the WSH effect to the next generation – wheat seeds had been purchased from a company and not exposed in any way to this water stress history. Collectively, these experiments demonstrate that drought legacy effects are likely transmitted through the seed microbiota, which are then capable of modulating plant responses to subsequent stress.

In 2016, bacterial seed endophyte richness tracked the rainfall gradient (100–75% > 50–25%), revealing an immediate internal loss of seed diversity. This early drought imprint was absent in fungal endophytes, pointing to a different response pattern of bacteria and fungi in short-term sensitivity to water stress inside seeds. These findings are in line with the previous reports that fungi can withstand better water limitation than bacteria (Manzoni, Schimel and Porporato 2012; Fuchslueger *et al*. 2014; Azarbad *et al*. 2018; Oram *et al*. 2025). Our results further revealed 2017 as a transient richness rebound inside seeds, where endophytic bacterial richness increased more under dry conditions (50–25% rainfall treatment) than respective treatment in 2016. Thus, drought-history seeds had recovered diversity internally, a feature specific to bacteria and not mirrored by fungal endophytes. We observed that across 2016 and 2017, the richness of bacterial endophytes correlated positively with yield, indicating that higher seed endophytic diversity resulted in greater productivity. Such a relationship aligns with ecological theory, predicting that diverse microbiota provide functional redundancy (Ramond, Galand and Logares 2025), which can, in turn, improve host stability under stress. This is consistent with experimental work demonstrating that soil microbial diversity buffers crop productivity under water limitation (Prudent *et al*. 2020). However, it is important to note that in our study, the strength of the diversity-yield relationship varied with rainfall regime and year in such a way that the effect of richness on yield was strongest under dry conditions in 2016 and under wet conditions in 2017, whereas the opposite combinations were non-significant. These results indicate that the positive influence of bacterial richness on yield depends on the rainfall treatment, with a year-specific pattern. The link between diversity and yield was further supported by the community composition analysis, where yield in 2017 was maximized when seed bacterial communities retained compositional features of the previous year’s drought-conditioned microbiota. Therefore, the compositional shift in seed bacterial community, together with changes in diversity, can be one of the main determinants of host productivity under stress factors.

Several of the persistent ASVs identified in this study belong to microbial taxa previously shown to promote plant growth, drought resilience, or disease suppression. *Pantoea* has been reported as a seed endophyte in wheat, enriched in drought-susceptible lines but not in tolerant cultivars (Hone *et al*. 2021); and in forage seeds, *Pantoea* isolates promoted germination under salt stress through IAA production and osmotic tolerance (Dai *et al*. 2020). In the case of fungi groups, some members of *Cladosporium* have been reported as one of the predominant fungal taxa in wheat seeds under drought stress conditions across generations (Vujanovic, Islam and Daida 2019), and *Alternaria* functions as a seed-borne endophyte that enhances wheat drought tolerance by producing auxin (IAA), stimulating antioxidant defenses and promoting osmotic adjustment (Qiang *et al*. 2019). Importantly, some of the taxa that persisted across generations in the transgenerational field experiment overlap with cultured, osmotolerant members that we previously isolated from the same experimental system, including a *Cladosporium* from roots and multiple bacterial isolates (*Bacillus* and *Pseudomonas*, including seed-derived *Bacillus*) that were able to increase wheat fresh biomass under drought after inoculation in the wheat plant environment (Agoussar *et al*. 2021; Agoussar, Tremblay and Yergeau 2025). Together, these lines of evidence support the interpretation that at least a subset of the taxa retained across generations in our field experiment are promising candidates for microbiome-mediated contributions to drought-associated plant performance.

Since the seed-associated microbiota is thought to be the preferred source of microorganisms for developing plants (Abdelfattah *et al*. 2021, 2023; Kim and Lee 2021; Moroenyane, Tremblay and Yergeau 2021) and that plant-associated microorganisms can impact the host phenotype (Li *et al*. 2019; Hone *et al*. 2021; Azarbad and Junker 2024), it would make sense that divergence in seed microbial communities would affect the performance of plants grown from these seeds. Indeed, the transgenerational field test demonstrated that the environmental history of the mother plants, experienced through rainfall manipulation, shaped the capacity of the daughter plants to tolerate drought. In 2018, AC Nass seeds originating from wetter parental plots (100% and 75% rainfall) exhibited yield declines under shelter (reduced precipitation), whereas those derived from dry parental plots (50% and 25%) maintained stable productivity. This cultivar- and origin-specific pattern suggests that drought-exposed parental environments can prime the next generation for improved performance under water limitation. The vertical transmission of microbiota through seeds is increasingly recognized as a key pathway for maintaining plant performance across generations. For example, a *Sphingomonas melonis* seed endophyte confers disease resistance in rice via a defined metabolite (anthranilic acid), antagonizing a pathogen’s virulence program, and is maintained transgenerationally in the seed endosphere (Matsumoto *et al*. 2021). In common bean lines (*Phaseolus vulgaris* L.), vertical transmission of seed endophytes was stable across three plant generations, even under drought and nutrient excess (the experiment was conducted in an indoor growth chamber). A conserved core of bacterial taxa, including *Pseudomonas*, *Bacillus*, *Massilia*, and *Sphingomonas*, persisted despite environmental stress, and most were absent from soil and rhizosphere compartments, indicating inheritance rather than reacquisition (Sulesky-Grieb *et al*. 2024). Vertical transmission has been shown from acorn embryo to seedling tissues in *Quercus robur*, with phyllosphere communities more closely reflecting the seed microbiota than root systems (Abdelfattah *et al*. 2021). Large-scale field work in potato indicated that seed origin imprints both microbiota and next-season performance (Song *et al*. 2025), and across spinach seed lots, higher relative abundances of *Massilia* was associated with damping-off suppression (Diakaki *et al*. 2025). Collectively, these results highlight that seed-borne communities are not passive, and their diversity and composition influence plant microbiota and plant performance in the next generation.

While the rainfall manipulation experiment and the transgenerational field test revealed that drought leaves a signature in seed-associated bacterial communities that correlates with yield stability, these results alone could not exclude the contribution of other non-microbial inheritance mechanisms. An example of such a mechanism is epigenetics, which consists of biochemical modifications to the chromosomes that do not involve a change in the nucleotide sequence, such as DNA methylation and histone modification, and that confer phenotypic plasticity to plants. This process plays a role in plant responses and adaptation to stress, including drought (Labra *et al*. 2002; Wang *et al*. 2011) and salinity (Zhong, Xu and Wang 2009). Interestingly, epigenetic modifications and the tolerance to abiotic stressors are often heritable in plants and can thus be passed on to subsequent generations (Boyko *et al*. 2007; Boyko and Kovalchuk 2008). However, the irrigation history greenhouse experiment provided another line of evidence pointing toward a microbial origin of the legacy effects observed. In this experiment, the seeds used in the greenhouse were collected from plants grown in soils that had experienced different long-term irrigation regimes but were never themselves exposed to those environmental histories other than through the soil microorganisms. Yet, the performance of the daughter plants emerging from those seeds varied according to the historical irrigation regime of the field of origin. When grown under controlled greenhouse drought treatments, plants derived from continuous WSH fields exhibited distinct physiological and microbial profiles compared with those from intermittent WSH fields. Specifically, AC Nass plants originating from seeds collected in the continuous WSH fields maintained higher photosynthetic assimilation and water-use efficiency under drought. These physiological advantages were paralleled by an increase in root bacterial diversity and a compositional shift, with drought-adapted plants harboring a distinct assemblage of taxa, including *Pseudoflavitalea*, *Bradyrhizobium*, and *Ralstonia*, also detected in their seed epiphyte pools. Because these seeds were produced in a single generation under identical climatic conditions, the observed differences must have been mediated through changes in the seed microbiota. Alternatively, it is possible that drought-conditioned microbiota indirectly modulated plant epigenetic states, as has been reported for other fungal and bacterial endophytes influencing host methylation patterns (Hubbard, Germida and Vujanovic 2014). However, even in such a case, the phenomenon would remain microbial in origin, with microorganisms serving as the drivers of transgenerational drought resilience. Another line of evidence strengthens a microbial interpretation for the legacy effects is metatranscriptomic evidence from the rainfall manipulation experiment where we previously showed that, as soil water content declines, the dominant transcriptomic shifts occur in the microbial partners (fungi in roots; bacteria in rhizosphere), whereas the plant contributes a smaller fraction of the differentially abundant transcripts (Pande *et al*. 2023).

Similar to the pattern observed in the rainfall manipulation experiment, in the irrigation history greenhouse experiment, the strength of microbial legacy effects was not uniform across wheat cultivars. Cultivar-dependent microbial assembly has been reported in multiple systems (Azarbad *et al*. 2020; Quiza *et al*. 2023; Michl *et al*. 2024). Crop breeding efforts have begun to recognize the potential of microbiota-assisted selection, proposing the integration of microbial inheritance and stress responsiveness as traits under selection (Cernava 2024). The predictive link between seed microbiota composition and the next generation performance under stress, observed in different plants (Dargiri and Samsampour 2025; Diakaki *et al*. 2025; Song *et al*. 2025), highlights the viability of this approach. As climate variability increases, the capacity of crops to buffer environmental extremes will depend not only on their genetic architecture but on the microbial partnerships they inherit.

Across different microbial systems, historical exposure to stress has been shown to consolidate resilience by enriching for specific taxa capable of sustaining community function under upcoming disturbance. For instance, it has been demonstrated that tree seedlings inoculated with soil microbial communities sourced from climate-matched sites survived better under the corresponding heat, cold, or drought stresses (Afkhami 2023; Allsup, George and Lankau 2023). In marine mesocosms, communities preconditioned to moderate stress (acidification) retain higher taxonomic and genomic diversity following severe disturbance (Xu *et al*. 2025), while in synthetic aquatic systems, prior disturbance (oxidative and dilution stressors) accelerates recovery trajectories and stabilizes reassembly around core taxa (Cairns *et al*. 2025). Similarly, evidence from our experiments demonstrates that seed microbiota retain a memory of drought, with legacy signals becoming especially pronounced under the driest environmental conditions. This suggests that microbial communities preconditioned by initial stress can confer enhanced adaptive value when host plants encounter comparable abiotic challenges. Such a context-dependent amplification of legacy effects aligns with findings in soil microbiota, where functional imprints of historical precipitation emerged most clearly during extreme water limitation (Broderick *et al*. 2025). Therefore, we suggest that the strength of microbial legacies is modulated not only by historical exposure but also by the severity of current stress.

In conclusion, our experiments in eastern and western Canada showed that the seed epiphytic and endophytic microbiota serve as ecological archives of the stress history of plants, and that seed bacterial and fungal communities retain and transmit drought history across plant generations. These insights open new avenues for microbiome-informed agriculture. Identifying, preserving, or reintroducing stress-adapted microbial consortia into seed systems could enhance crop resilience under water limitation without the need for genetic modification or chemical inputs.

## Material and Methods

### Experiment 1: Rainfall manipulation experiment (Laval, Quebec, Canada)

#### The effects of rainfall manipulation on wheat yields and wheat-associated microbiota

In 2016, we designed a multi-year field experiment at the Armand-Frappier Santé Biotechnologie Research Center campus (Laval, QC, Canada). The field (23 m × 17 m) was established based on a randomized complete block design, which was divided into six blocks where blocks were separated by 2 m from each other (Fig. 1a). Within each block, eight plots (4 m^2^) were arranged, with 1 m spacing between adjacent plots. Each of the eight plots in a block was randomly assigned to one of four rainfall manipulation treatments: 100% precipitation (control), 75% precipitation, 50% precipitation, or 25% precipitation (Wang *et al*. 2022; Asad *et al*. 2023). From May to October each year, all plots except the control were partially covered with wooden shelters that supported UV-transparent plastic sheets. Rainfall intercepted by these sheets was funneled into gutters and 20-L collection buckets, which were manually emptied after significant rainfall events. Two wheat cultivars, drought-tolerant (*Triticum turgidum subsp. durum* cv. Strongfield) and sensitive (*Triticum aestivum* cv. AC Nass), were grown in the field. This resulted in a total of 48 plots (4 precipitations × 2 cultivars × 6 replicate blocks). For the first year (2016), the seeds were obtained from Agriculture and Agri-Food Canada. In subsequent years (2017 and 2018), seeds were harvested at the end of the growing season from each plot and used to reseed the same plots in May of the following year. In each plot, approximately 2,000 seeds were distributed in ten rows, resulting in a density of 500 seeds/m^²^. In terms of field management, to ensure that only the target wheat cultivars were grown in each plot, the experimental field was manually weeded at biweekly intervals throughout the growing season each year. No herbicides, insecticides, or other agrochemicals were applied. Seeds were collected at the end of August each year, resulting in 3 different plant generations (2016, 2017, and 2018: Fig. 1b), and epiphytes and endophyte seed microbiota were isolated after each harvest. Although seeds were collected in 2018, the yield data from that year were unfortunately not retained due to an unexpected data loss and could not be included in the final statistical analysis.

Soil samples were taken before seeding (T0) and from the rhizosphere, roots, and leaves on July 20th during each growing season. They were then immediately stored at -20 °C in the laboratory prior to DNA extraction. Monthly precipitation data for May through August in 2016, 2017, and 2018 were obtained from the Montreal International Airport (QC) weather station located 8.5 km from the experimental site. These records were used to track seasonal rainfall trends and interannual climatic variability (Fig. 1c). To complement these data and quantify soil moisture conditions under each rainfall treatment at the time of the sampling, soil samples were collected from each plot and used to determine gravimetric soil water content (Fig. 1d). For this, fresh soil was weighed, oven-dried at 75°C for 48 hours, and then reweighed to calculate the water content as the difference between the fresh and dry weights relative to the dry weight.

### Experiment 2: Transgenerational field experiment: testing pre-adaptation across generations (Laval, Quebec, Canada)

In 2017, a second field experiment was established to test whether seeds collected under the different precipitation regimes of Experiment 1 in 2016 were pre-adapted to reduced water availability. The field dedicated to this experiment was established next to experiment 1. Seeds were harvested from the main experiment at the end of the 2016 growing season from plots that had received one of four precipitation levels (100%, 75%, 50%, 25%) and belonged to either the drought-tolerant (Strongfield) or drought-sensitive (AC Nass) cultivar. This resulted in eight distinct seed origins (4 precipitation histories × 2 cultivars).

The field was divided into two blocks. Each block contained six plots, designated for one of two main conditions: (A) with shelter (reduced precipitation: 25% of the ambient precipitation) and (B) without shelter (ambient precipitation: receiving natural rainfall). Within these two blocks, every plot was randomly assigned to one of the six blocks from Experiment 1 (Fig. S6). Within each plot, the area was subdivided into eight subplots. Each subplot was dedicated to one of the eight seed origins. Thus, each subplot was planted with seeds from one of the eight seed origins, ensuring that every combination of seed history and precipitation condition was evaluated across multiple replicates. This factorial design (2 precipitation treatments × 8 seed origins × 2 blocks) allowed us to examine whether wheat seeds originated from a specific water regime in 2016 exhibited differential yield responses when subjected to new water conditions in 2017. The experiment was repeated in 2018 using seeds harvested from the 2017 experiment. Seeds were sown in May 2017 and 2018 using density and row spacing similar to those in Experiment 1.

### Experiment 3: Irrigation history greenhouse experiment (Swift Current, Saskatchewan, Canada)

To evaluate whether the seed microbiota-mediated legacy effects under drought are consistent across distinct agroecosystems, we extended our investigation beyond the original Laval field system in eastern Canada to a geographically and climatically distinct agricultural region in western Canada (Fig. 5a). For that, we took advantage of a field experiment in Swift Current, Saskatchewan (Azarbad *et al*. 2022). A field experiment was conducted in two adjacent agricultural fields differing in historical irrigation regimes: one field had received no supplemental irrigation for 37 years (continuous water stress history, WSH), while the other was regularly irrigated (intermittent WSH). Four wheat cultivars, including AC Nass and Strongfield, were grown under two contemporary irrigation conditions (irrigated and non-irrigated). This design allowed disentangling the effects of historical and contemporary water stress on plant traits and their associated bacteria and fungi communities (Azarbad *et al*. 2022). This experimental site differs from the Laval experiment in terms of precipitation patterns, temperature regime, and long-term management history. This provides an ideal setting to test whether seed microbiota dynamics and transgenerational plant responses observed under Laval experimental conditions in the eastern part of Canada are conserved or context-dependent across other environmental settings. Seeds were sampled at harvest from each treatment plot and analyzed for their endophytic and epiphytic microbial communities. We have previously shown that seed epiphytic bacterial and fungal communities were shaped by the historical water stress regime, suggesting strong microbial legacy effects that were imprinted on the seed microbiota (Azarbad *et al*. 2022).

Seeds harvested from the field Swift Current experiment were grown in a greenhouse to assess their growth under water-limited conditions (Fig. 5b). The greenhouse experiment was carried out in 2019, where 8 seeds from each field plot were placed in pots (1 L flat-bottom), filled with sterilized potting soil (800 g of dry weight soil). Pots in the greenhouse were completely randomized in four blocks and were regularly rotated on the greenhouse bench to avoid any spatial/greenhouse effects. After four weeks of growth under 50% soil water holding capacity (SWHC), the soil water content in half the pots was adjusted to 20% SWHC. This resulted in 120 pots (60 batches of seeds harvested from the field experiment × 2 SWHC). Soil water content was maintained at either 20% or 50% of the SWHC by weighing the pots every second day at a regular time (between 7:30-9:30 am). Following four weeks of exposure to 50% and 20% SWHC, photosynthetic performance was assessed using a CIRAS-3 portable infrared gas analyzer (PP Systems, USA), which allows simultaneous quantification of multiple leaf-level gas exchange parameters. Among these, net assimilation rate (A; μmol CO₂ m⁻² s⁻¹) and photosynthetic water use efficiency (WUE; mmol CO₂ mol⁻¹ H₂O) were selected, as they are physiologically informative metrics that reflect carbon fixation capacity and the balance between carbon gain and transpirational water loss, respectively. Measurements were conducted on five randomly selected plants per pot. For consistency and to minimize intra-plant variation, all measurements were taken on the same fully expanded leaf per plant. Following gas exchange measurements, the plants were assessed for aboveground plant height (cm), measured from the base to the tip of the longest shoot. Plants were then harvested, rhizosphere samples were collected for DNA extraction and microbial sequencing, and their shoots were dried at 75 °C to a constant weight to determine the aboveground dry biomass.

### Extracting the seed endophytes and epiphytes

Detailed information on seed epiphytes and endophytes DNA extraction is presented in (Azarbad *et al*. 2022). For epiphytes, 10 g of seeds from each plot were shaken for 1 h in sterile peptone buffer (Kim *et al*. 2006), and cell pellets obtained after centrifugation were resuspended in TEP buffer and stored at −20 °C until DNA extraction. For endophytes, the same seed material was surface-sterilized (95% ethanol, sodium hypochlorite, repeated sterile water rinses) following (Sun *et al*. 2008) (with some modifications), and sterility was confirmed by plating the final rinse on LB and TSB agar (Fig. S7). Sterilized seeds were flash-frozen in liquid nitrogen, ground with a mortar and pestle, and 0.5 g aliquots were stored at −20 °C for subsequent DNA extraction.

### DNA extraction, amplicon library preparation, and sequencing

For DNA extraction, leaf and root samples were ground in powder using liquid nitrogen with a mortar and pestle. DNA was extracted from 0.5 g of bulk soil, rhizosphere, roots, leaves and the seed (endophyte) samples using a phenol-chloroform extraction method (Dellaporta, Wood and Hicks 1983). For seed epiphyte extraction, pellets which were resuspended in 1000 μl of TEP buffer were used directly for phenol-chloroform DNA extraction. Detailed information on DNA extraction is presented in Azarbad *et al*. (2018). Sequencing libraries were prepared using two steps approach as described previously (Yergeau *et al*. 2015). Amplicon libraries were prepared for the bacterial 16S rRNA gene using the universal primers 520F and 799R targeting the V4 region (Edwards *et al*. 2007) and for the fungal ITS1 region using ITS1F and 58A2R primers (Martin and Rygiewicz 2005). Samples corresponding to the bacteria and fungi were pooled separately and sequenced on an Illumina MiSeq sequencer (250PE) at the Centre d’expertise et de séquençage Genome Québec (Montréal, Canada). For the Laval experiment, to minimize technical variability, all seed epiphyte and endophyte samples were submitted together and processed simultaneously for sequencing, ensuring identical library preparation and sequencing runs.

### Analysis of sequencing data

Sequences were analyzed using AmpliconTagger (Tremblay and Yergeau 2019). Briefly, raw reads were scanned for sequencing adapters and PhiX spike-in sequences. The remaining reads were filtered based on quality (Phred) score and remaining sequences were dereplicated/clustered at 100% identity and then processed for generating Amplicon Sequence Variants (ASVs) (DADA2 v1.12.1) (PMID:27214047). Chimeras were removed with DADA2’s internal removeBimeraDeNovo (method = “consensus”) method followed by UCHIME reference (Edgar *et al*. 2011). ASVs for which abundance across all samples were lower than 3 were discarded. ASVs were assigned a taxonomic lineage with the RDP classifier (PMID: 17586664) using an in-house training set containing the complete Silva release 128 database (PMID:23193283) supplemented with eukaryotic sequences from the Silva database and a customized set of mitochondria, plasmid and bacterial 16S sequences. For ITS ASVs, a training set containing the Unite DB was used (sh_general_release_s_04.02.2020 version). The RDP classifier assigns a score (0 to 1) to each taxonomic depth of each ASV. Each taxonomic depth having a score ≥ 0.5 were kept to reconstruct the final lineage. Taxonomic lineages were combined with the cluster abundance matrix obtained above to generate a raw ASV table, from which a bacterial organisms ASV table was generated. Taxonomic summaries were computed using the QIIME v1.9.1 software suite (PMID: 20383131, 22161565) using the ASV table of each data type.

### Statistical analyses

Statistical analyses were performed in R Studio (v 4.3.2, The R Foundation for Statistical Computing). For the rainfall manipulation experiment (Experiment 1), the variation in soil water content was assessed by ANOVA, using the *aov*() function, with rainfall treatment (100, 75, 50, 25% of ambient) and year (2016–2018) as fixed factors (and block as a random effect), and the interaction between them. For yield, ANOVA analyses were performed separately for AC Nass and Strongfield for 2016 and 2017 to account for cultivar and year-specific responses. When ANOVA indicated significant effects, pairwise differences among treatments were identified using Tukey’s HSD post hoc test (*emmeans* package). To verify the possible direct and interactive effects of rainfall treatment (100, 75, 50, 25%), generation (2016–2018), cultivar (AC Nass, Strongfield), and block as a random factor on rhizosphere, root, and leaf, seed epiphytes, and endophytes bacterial and fungal alpha diversity (observed richness), Kruskal-Wallis tests were performed. The effect of experimental factors on rhizosphere, root, and leaf, seed epiphytes, and endophytes, and bacterial and fungal community composition was visualized using principal coordinate analyses (PCoA). The impact of the experimental parameters and their interactions on bacterial and fungal community composition was tested using Permanova with 1000 permutations (adonis function from the vegan package in R). The Pemanovas and the PCoAs were performed on relative abundance (using Bray–Curtis dissimilarity). ANOVA was performed to determine the impact of each of the experimental factors and their interactions on the relative abundance of dominant bacterial and fungal genera associated with seed epiphytes and endophytes.

Further analyses focused on testing whether seed bacterial endophyte and fungal epiphytes diversity and community composition predicted variation in wheat yield under different irrigation regimes and across growing years (2016–2017). Rainfall levels were grouped into: dry (25 % + 50 %) and wet (75 % + 100 %) rainfall regimes. Differences in seed bacterial endophyte richness between rainfall regimes (wet, dry) and years (2016–2017) were evaluated using linear model. Estimated marginal means (EMMs) and their 95 % confidence limits were extracted using the *emmeans* package to visualize group differences. The association between richness and yield was first assessed using Spearman’s rank correlations within each year. To evaluate the effect of richness while accounting for covariates, we fitted a linear mixed-effects model of yield as a function of richness, rainfall regime (wet/dry), and year, including soil water content (SWC) as a covariate and block as a random intercept using the *lmer()* function. Richness and soil water content (SWC) were standardized (z-scores) prior to modeling so that estimated slopes (β) represent the expected change in yield per one-standard-deviation increase in each predictor. From this model, marginal slopes (β ±95% confidence intervals) were extracted using the *emtrends()* function to estimate the effect of richness within each rainfall regime and year. P-values were adjusted for multiple testing using the Benjamini–Hochberg false discovery rate (FDR ≤ 0.05). To test whether seed community composition in 2017 reflected a compositional “legacy” of the 2016 drought microbiota, a centroid representing the mean relative-abundance profile of the 2016–dry group was calculated, and each 2017 community’s distance to this centroid (DistTo2016Dry) was used as a quantitative measure of dissimilarity. Yield was then modeled as a function of compositional distance, rainfall origin, and their interaction, controlling for SWC and block effects. Model output provided the main and interaction effects as well as per rain-group slopes (β ± 95 % CI). In addition, Spearman’s ρ between yield and distance was computed for each rainfall group to quantify the strength and direction of association.

For the transgenerational field test (Experiment 2), yield data from the 2017–2018 rain-out validation were analysed separately for AC Nass and Strongfield. For each cultivar, we tested the effect of shelter (with vs. without) within each seed precipitation history (100, 75, 50, 25%) using two-sample t-tests. This allowed us to assess whether seeds originating from wetter or drier rainfall histories responded differently to shelter-imposed drought in the subsequent generation.

In the irrigation history greenhouse experiment, for physiological (net assimilation rate and water use efficiency) and morphological traits (shoot biomass and plant height), ANOVA tests were performed with soil water holding capacity (SWHC; 50% vs. 20%), field history (with vs. without water stress history), irrigation regime (irrigated vs. non-irrigated), and cultivar (AC Nass vs. Strongfield) as fixed factors. For the root bacterial alpha diversity, effects of field history, irrigation, cultivar, and SWHC were tested using ANOVA, and significant interactions were reported. To identify shared ASVs across seed and root compartments, a set of 29 core ASVs was compiled and visualized using a heatmap (*pheatmap* package). For that, relative abundances were log-transformed, and clustering was based on Euclidean distance.

## Data availability

The data that support the findings of this study are available in the supplementary material of this article. The raw amplicon data sets and associated metadata related to the Laval experiment are available through NCBI BioProject accession PRJNA767855. The data associated with the Saskatchewan experiment are available in the NCBI Sequence Read Archive (SRA) under the BioProject accession PRJNA736197.

## Acknowledgments

We would like to thank all the members, past and current, of the mECO:LABS for their help in setting up and maintaining the field experiment, in particular Usman Irshad, Sara Correa-Garcia, Itumeleng Moroenyane, Simon Gallichand Desmeules, Liliana Quiza, Lilia Bouyoucef, Éloïse Adam-Granger, Deanna Chinnerman, and Karelle Rheault. We thank Mathieu Gaudreault for his excellent technical support. We would also like to thank the Cereal Breeding research group at the Swift Current Research and Development Centre of Agriculture and Agri-food Canada for their help with the Swift Current field experiment. We would also like to thank Derry Wallis and Bianca Evans for their assistance in setting up, maintaining, and harvesting the greenhouse experiment. This work was funded by the Natural Sciences and Engineering Research Council of Canada (Discovery grant RGPIN-2014-05274 and Strategic grant for projects STPGP 494702 to EY). HA was supported by a Fonds de recherche du Québec-Nature et technologies (FRQNT) and a Fondation Armand-Frappier postdoctoral fellowships. We also wish to acknowledge Compute Canada for access to the University of Waterloo’s High-Performance Computing (HPC) infrastructure (Graham system) through a resources allocation granted to EY.

## Author contributions

HA and EY designed the study. PP, CGL, and JAD made substantial contributions to maintain the Laval field experiments. JAD contributed to DNA extraction and library preparations. AA contributed to the seed DNA extractions. LB planned and performed the Swift Current field and greenhouse experiment. JT performed the bioinformatic analyses and bioinformatic methods writing. HA analyzed the data and wrote the manuscript. All co-authors participated in reviewing and editing the final text. The authors declare no competing interests.

