## Supplementary for "Seed microbiota legacy mitigates the effect of drought in wheat"

**Table S1.** Kruskal-Wallis test for the effects of treatment (T), cultivars (C), generations (G), block (B), and their interactions on bacteria richness diversity of bulk soil, rhizosphere, leaf, root, seed epiphytes, and endophytes communities.

| | **Bacteria – Richness (including cultivar as factor)** | | | | | | | | | | | | | | |
|---|---|---|---|---|---|---|---|---|---|---|---|---|---|---|---|
| | Rhizosphere | | | Root | | | Leaf | | | Seed Epiphytes | | | Seed Endophytes | | |
| | chi-squared | df | p-value | chi-squared | df | p-value | chi-squared | df | p-value | chi-squared | df | p-value | chi-squared | df | p-value |
| T | 1.385 | 3 | 0.709 | 3.502 | 3 | 0.321 | 0.405 | 3 | 0.939 | 1.008 | 3 | 0.799 | 4.159 | 3 | 0.245 |
| C | 0.314 | 1 | 0.576 | 2.468 | 1 | 0.116 | 0.147 | 1 | 0.701 | 2.730 | 1 | 0.098 | 7.371 | 1 | **0.007** |
| G | 8.482 | 2 | **0.014** | 0.013 | 2 | 0.994 | 22.791 | 2 | **0.000** | 20.074 | 2 | 0.000 | 11.642 | 2 | **0.003** |
| B | 3.945 | 5 | 0.557 | 10.237 | 5 | 0.069 | 1.956 | 5 | 0.855 | 5.468 | 5 | 0.361 | 8.941 | 5 | 0.111 |
| T × C | 4.892 | 7 | 0.673 | 9.162 | 7 | 0.241 | 3.156 | 7 | 0.870 | 4.416 | 7 | 0.731 | 13.491 | 7 | 0.061 |
| T × G | 19.064 | 11 | 0.060 | 9.707 | 11 | 0.557 | 26.886 | 11 | **0.005** | 31.469 | 11 | **0.001** | 19.437 | 11 | **0.054** |
| C × G | 9.639 | 4 | **0.047** | 3.200 | 4 | 0.525 | 36.335 | 4 | **0.000** | 20.352 | 4 | **0.000** | 22.814 | 4 | **0.000** |
| T× C × G | 25.492 | 19 | 0.145 | 16.050 | 19 | 0.654 | 44.943 | 19 | **0.001** | 33.674 | 19 | **0.020** | 35.148 | 19 | **0.013** |

| | **AC Nass** | | | | | | | | | | | | | | |
|---|---|---|---|---|---|---|---|---|---|---|---|---|---|---|---|
| | Rhizosphere | | | Root | | | Leaf | | | Seed Epiphytes | | | Seed Endophytes | | |
| | chi-squared | df | p-value | chi-squared | df | p-value | chi-squared | df | p-value | chi-squared | df | p-value | chi-squared | df | p-value |
| T | 2.422 | 3 | 0.490 | 5.039 | 3 | 0.169 | 0.849 | 3 | 0.838 | 1.012 | 3 | 0.798 | 3.474 | 3 | 0.324 |
| G | 9.596 | 2 | **0.008** | 0.715 | 2 | 0.700 | 32.642 | 2 | **0.000** | 18.363 | 2 | **0.000** | 13.968 | 2 | **0.001** |
| B | 2.513 | 5 | 0.775 | 10.215 | 5 | 0.069 | 5.471 | 5 | 0.361 | 2.820 | 5 | 0.728 | 3.997 | 5 | 0.550 |
| T × G | 14.436 | 11 | 0.210 | 11.097 | 11 | 0.435 | 37.283 | 11 | **0.000** | 31.160 | 11 | **0.001** | 23.000 | 11 | **0.018** |

| | **Strongfield** | | | | | | | | | | | | | | |
|---|---|---|---|---|---|---|---|---|---|---|---|---|---|---|---|
| | chi-squared | df | p-value | chi-squared | df | p-value | chi-squared | df | p-value | chi-squared | df | p-value | chi-squared | df | p-value |
| T | 2.633 | 3 | 0.452 | 2.397 | 3 | 0.494 | 2.940 | 3 | 0.401 | 0.685 | 3 | 0.877 | 2.410 | 3 | 0.492 |
| G | 0.055 | 1 | 0.815 | 0.045 | 1 | 0.831 | 0.495 | 1 | 0.482 | 0.943 | 1 | 0.331 | 1.688 | 1 | 0.194 |
| B | 2.672 | 5 | 0.750 | 4.758 | 5 | 0.446 | 9.535 | 5 | 0.090 | 10.203 | 5 | 0.070 | 8.532 | 5 | 0.129 |
| T × G | 11.402 | 7 | 0.122 | 3.331 | 7 | 0.853 | 4.632 | 7 | 0.705 | 2.494 | 7 | 0.928 | 4.613 | 7 | 0.707 |

**Table S2.** Kruskal-Wallis test for the effects of treatment (T), cultivars (C), generations (G), block (B), and their interactions on fungi richness diversity of bulk soil, rhizosphere, leaf, root, seed epiphytes, and endophytes communities.

| | **Fungi – Richness (including cultivar as factor)** | | | | | | | | | | | | | | |
|---|---|---|---|---|---|---|---|---|---|---|---|---|---|---|---|
| | Rhizosphere | | | Root | | | Leaf | | | Seed Epiphytes | | | Seed Endophytes | | |
| | chi-squared | df | p-value | chi-squared | df | p-value | chi-squared | df | p-value | chi-squared | df | p-value | chi-squared | df | p-value |
| T | 1.071 | 3 | 0.784 | 5.057 | 3 | 0.168 | 0.471 | 3 | 0.925 | 6.203 | 3 | 0.102 | 2.065 | 3 | 0.559 |
| C | 7.050 | 1 | **0.008** | 2.061 | 1 | 0.151 | 5.089 | 1 | 0.024 | 0.127 | 1 | 0.722 | 4.901 | 1 | **0.027** |
| G | 83.179 | 2 | **0.000** | 79.148 | 2 | **0.000** | 32.837 | 2 | **0.000** | 11.578 | 2 | **0.003** | 3.550 | 2 | 0.169 |
| B | 2.000 | 5 | 0.849 | 2.084 | 5 | 0.837 | 11.662 | 5 | **0.040** | 1.784 | 5 | 0.878 | 7.335 | 5 | 0.197 |
| T × C | 10.775 | 7 | 0.149 | 7.291 | 7 | 0.399 | 8.038 | 7 | 0.329 | 8.484 | 7 | 0.292 | 7.757 | 7 | 0.354 |
| T × G | 86.973 | 11 | **0.000** | 86.995 | 11 | **0.000** | 37.102 | 11 | **0.000** | 25.675 | 11 | **0.007** | 16.533 | 11 | 0.122 |
| C × G | 84.439 | 4 | **0.000** | 79.750 | 4 | **0.000** | 42.972 | 4 | **0.000** | 13.727 | 4 | **0.008** | 11.166 | 4 | **0.025** |
| T× C × G | 89.471 | 19 | **0.000** | 88.860 | 19 | **0.000** | 51.087 | 19 | **0.000** | 31.775 | 19 | **0.033** | 28.009 | 19 | 0.083 |

| | **AC Nass** | | | | | | | | | | | | | | |
|---|---|---|---|---|---|---|---|---|---|---|---|---|---|---|---|
| | Rhizosphere | | | Root | | | Leaf | | | Seed Epiphytes | | | Seed Endophytes | | |
| | chi-squared | df | p-value | chi-squared | df | p-value | chi-squared | df | p-value | chi-squared | df | p-value | chi-squared | df | p-value |
| T | 2.622 | 3 | 0.454 | 3.851 | 3 | 0.278 | 0.526 | 3 | 0.913 | 3.897 | 3 | 0.273 | 3.002 | 3 | 0.391 |
| G | 46.109 | 2 | **0.000** | 41.997 | 2 | **0.000** | 14.161 | 2 | **0.001** | 8.405 | 2 | **0.015** | 0.058 | 2 | 0.971 |
| B | 3.171 | 5 | 0.674 | 1.268 | 5 | 0.938 | 18.614 | 5 | **0.002** | 1.866 | 5 | 0.867 | 4.073 | 5 | 0.539 |
| T × G | 49.502 | 11 | **0.000** | 50.002 | 11 | **0.000** | 18.585 | 11 | 0.069 | 22.281 | 11 | **0.022** | 12.416 | 11 | 0.333 |

| | **Strongfiled** | | | | | | | | | | | | | | |
|---|---|---|---|---|---|---|---|---|---|---|---|---|---|---|---|
| | chi-squared | df | p-value | chi-squared | df | p-value | chi-squared | df | p-value | chi-squared | df | p-value | chi-squared | df | p-value |
| T | 1.829 | 3 | 0.609 | 2.397 | 3 | 0.479 | 3.370 | 3 | 0.338 | 3.669 | 3 | 0.300 | 0.472 | 3 | 0.925 |
| G | 32.772 | 1 | **0.000** | 0.045 | 1 | **0.000** | 22.120 | 1 | **0.000** | 5.731 | 1 | **0.017** | 5.674 | 1 | 0.017 |
| B | 1.292 | 5 | 0.936 | 4.758 | 5 | 0.873 | 0.683 | 5 | 0.984 | 1.096 | 5 | 0.954 | 4.097 | 5 | 0.536 |
| T × G | 35.850 | 7 | **0.000** | 3.331 | 7 | **0.000** | 26.818 | 7 | **0.000** | 9.646 | 7 | 0.210 | 10.269 | 7 | 0.174 |

**Table S3.** ANOVA tests for the effects of treatment (T), cultivars (C), generations (G), block (B) and their interactions on bulk soil, rhizosphere, leaf, root, seed epiphytes, and endophytes associated bacterial communities.

| | **Bacteria (including cultivar as factor)** | | | | | | | | | | | | | | |
|---|---|---|---|---|---|---|---|---|---|---|---|---|---|---|---|
| | Rhizosphere | | | Root | | | Leaf | | | Seed Epiphytes | | | Seed Endophytes | | |
| | F-value | df | p-value | F-value | df | p-value | F-value | df | p-value | F-value | df | p-value | F-value | df | p-value |
| T | 1.016 | 3 | 0.372 | 1.323 | 3 | **0.048** | 1.024 | 3 | 0.343 | 1.532 | 3 | **0.053** | 1.423 | 3 | 0.073 |
| C | 2.613 | 1 | **0.017** | 2.930 | 1 | **0.002** | 5.365 | 1 | **0.001** | 22.763 | 1 | **0.001** | 12.697 | 1 | **0.001** |
| G | 29.722 | 2 | **0.001** | 11.868 | 2 | **0.001** | 20.297 | 2 | **0.001** | 21.709 | 2 | **0.001** | 9.358 | 2 | **0.001** |
| B | 2.776 | 1 | **0.013** | 2.576 | 1 | **0.001** | 2.781 | 1 | **0.008** | 3.310 | 1 | **0.004** | 1.162 | 1 | 0.313 |
| T × C | 1.145 | 3 | 0.233 | 1.248 | 3 | 0.086 | 1.330 | 3 | 0.097 | 1.211 | 3 | 0.213 | 0.703 | 3 | 0.883 |
| T × G | 0.873 | 6 | 0.708 | 0.919 | 6 | 0.736 | 1.002 | 6 | 0.456 | 0.973 | 6 | 0.507 | 1.222 | 6 | 0.143 |
| C × G | 0.813 | 1 | 0.538 | 2.148 | 1 | **0.004** | 5.970 | 1 | **0.001** | 3.234 | 1 | **0.005** | 1.718 | 1 | 0.061 |
| T× C × G | 0.892 | 3 | 0.604 | 0.791 | 3 | 0.914 | 0.973 | 3 | 0.458 | 1.078 | 3 | 0.323 | 1.179 | 3 | 0.230 |

| | **Bacteria – AC Nass** | | | | | | | | | | | | | | |
|---|---|---|---|---|---|---|---|---|---|---|---|---|---|---|---|
| | Rhizosphere | | | Root | | | Leaf | | | Seed Epiphytes | | | Seed Endophytes | | |
| | F-value | df | p-value | F-value | df | p-value | F-value | df | p-value | F-value | df | p-value | F-value | df | p-value |
| T | 1.123 | 3 | 0.256 | 1.420 | 3 | **0.023** | 1.184 | 3 | 0.230 | 1.034 | 3 | 0.387 | 1.265 | 3 | 0.186 |
| G | 19.044 | 2 | **0.001** | 9.401 | 2 | **0.001** | 19.352 | 2 | **0.001** | 21.017 | 2 | **0.001** | 8.938 | 2 | **0.001** |
| B | 2.028 | 5 | **0.048** | 2.269 | 5 | **0.001** | 1.942 | 5 | 0.076 | 2.652 | 5 | **0.024** | 0.902 | 5 | 0.527 |
| T × G | 0.789 | 6 | 0.868 | 0.957 | 6 | 0.564 | 0.967 | 6 | 0.515 | 1.124 | 6 | 0.276 | 1.358 | 6 | **0.042** |

| | **Bacteria – Strongfield** | | | | | | | | | | | | | | |
|---|---|---|---|---|---|---|---|---|---|---|---|---|---|---|---|
| | F-value | df | p-value | F-value | df | p-value | F-value | df | p-value | F-value | df | p-value | F-value | df | p-value |
| T | 1.038 | 3 | 0.360 | 1.287 | 3 | 0.125 | 1.230 | 3 | 0.145 | 1.447 | 3 | 0.071 | 0.841 | 3 | 0.722 |
| G | 22.215 | 1 | **0.001** | 9.277 | 1 | **0.001** | 8.853 | 1 | **0.001** | 8.545 | 1 | **0.001** | 3.283 | 1 | **0.001** |
| B | 1.640 | 5 | 0.087 | 1.480 | 5 | **0.009** | 1.324 | 5 | 0.066 | 2.443 | 5 | **0.012** | 1.441 | 5 | 0.151 |
| T × G | 1.054 | 3 | 0.345 | 0.954 | 3 | 0.709 | 1.011 | 3 | 0.404 | 0.826 | 3 | 0.712 | 0.931 | 3 | 0.423 |

**Table S4.** ANOVA tests for the effects of treatment (T), cultivars (C), generations (G), block (B) and their interactions on bulk soil, rhizosphere, leaf, root, seed epiphytes, and endophytes associated fungi communities.

| | **Fungi (including cultivar as factor)** | | | | | | | | | | | | | | |
|---|---|---|---|---|---|---|---|---|---|---|---|---|---|---|---|
| | Rhizosphere | | | Root | | | Leaf | | | Seed Epiphytes | | | Seed Endophytes | | |
| | F-value | df | p-value | F-value | df | p-value | F-value | df | p-value | F-value | df | p-value | F-value | df | p-value |
| T | 0.790 | 3 | 0.782 | 1.506 | 3 | **0.022** | 1.407 | 3 | 0.168 | 1.656 | 3 | 0.060 | 0.856 | 3 | 0.514 |
| C | 2.140 | 1 | **0.032** | 3.230 | 1 | **0.003** | 28.628 | 1 | **0.001** | 27.504 | 1 | **0.001** | 35.580 | 1 | **0.001** |
| G | 23.805 | 2 | **0.001** | 12.313 | 2 | **0.001** | 13.064 | 2 | **0.001** | 49.065 | 2 | **0.001** | 29.685 | 2 | **0.001** |
| B | 2.288 | 1 | **0.019** | 2.700 | 1 | **0.004** | 10.666 | 1 | **0.001** | 3.225 | 1 | **0.009** | 1.163 | 1 | 0.278 |
| T × C | 1.189 | 3 | 0.215 | 0.812 | 3 | 0.809 | 0.488 | 3 | 0.925 | 1.934 | 3 | **0.023** | 1.628 | 3 | 0.093 |
| T × G | 0.895 | 6 | 0.689 | 1.024 | 6 | 0.372 | 1.643 | 6 | **0.037** | 1.048 | 6 | 0.398 | 0.958 | 6 | 0.500 |
| C × G | 0.741 | 1 | 0.680 | 1.851 | 1 | **0.042** | 10.232 | 1 | **0.001** | 5.689 | 1 | **0.00**1 | 3.322 | 1 | **0.018** |
| T× C × G | 1.166 | 3 | 0.235 | 0.983 | 3 | 0.503 | 0.666 | 3 | 0.720 | 1.189 | 3 | 0.247 | 0.764 | 3 | 0.649 |

| | **Fungi – AC Nass** | | | | | | | | | | | | | | |
|---|---|---|---|---|---|---|---|---|---|---|---|---|---|---|---|
| | Rhizosphere | | | Root | | | Leaf | | | Seed Epiphytes | | | Seed Endophytes | | |
| | F-value | df | p-value | F-value | df | p-value | F-value | df | p-value | F-value | df | p-value | F-value | df | p-value |
| T | 0.927 | 3 | 0.567 | 1.276 | 3 | 0.076 | 1.073 | 3 | 0.353 | 1.809 | 3 | **0.049** | 1.004 | 3 | 0.436 |
| G | 15.943 | 2 | **0.001** | 7.839 | 2 | **0.001** | 13.305 | 2 | **0.001** | 50.055 | 2 | **0.001** | 25.594 | 2 | **0.001** |
| B | 1.394 | 5 | **0.049** | 1.366 | 5 | **0.020** | 3.232 | 5 | **0.002** | 2.373 | 5 | **0.003** | 1.048 | 5 | 0.402 |
| T × G | 0.928 | 6 | 0.606 | 1.178 | 6 | 0.110 | 1.264 | 6 | 0.186 | 1.411 | 6 | 0.124 | 1.118 | 6 | 0.319 |
| | **Fungi – Strongfield** | | | | | | | | | | | | | | |
| | F-value | df | p-value | F-value | df | p-value | F-value | df | p-value | F-value | df | p-value | F-value | df | p-value |
| T | 1.211 | 3 | 0.219 | 1.096 | 3 | 0.274 | 0.978 | 3 | 0.471 | 1.829 | 3 | **0.035** | 1.324 | 3 | 0.218 |
| G | 17.832 | 1 | **0.001** | 11.075 | 1 | **0.001** | 5.714 | 1 | **0.001** | 23.130 | 1 | **0.001** | 15.448 | 1 | **0.001** |
| B | 1.594 | 5 | **0.042** | 1.204 | 5 | 0.164 | 0.803 | 5 | 0.757 | 1.358 | 5 | 0.138 | 1.082 | 5 | 0.373 |
| T × G | 1.223 | 3 | 0.216 | 0.628 | 3 | 0.973 | 1.489 | 3 | 0.119 | 1.025 | 3 | 0.378 | 0.641 | 3 | 0.806 |

**Table S5.** ANOVA tests for the effects of SWHC, field history, cultivar, irrigation, and their interactions on plant physiological (net assimilation rate and water use efficiency) and morphological traits (shoot biomassone health and plant height).

| **The greenhouse experiment** | | | | | | | | |
|---|---|---|---|---|---|---|---|---|
| **Factor** | Shoot biomass | | Plant height | | Assimilation Rate | | WUE | |
| | F | P | F | P | F | P | F | P |
| SWHC (%) | 56.236 | **0.000** | 83.07 | **0.000** | 38.954 | **0.000** | 150.5 | **0.000** |
| Field history | 0.179 | 0.675 | 0.601 | 0.442 | 3.625 | 0.058 | 1.748 | 0.187 |
| Cultivar | 36.730 | **0.000** | 75.966 | **0.000** | 23.077 | **0.000** | 9.002 | 0.003 |
| Irrigation | 0.009 | 0.925 | 0.116 | 0.735 | 0.563 | 0.454 | 1.343 | 0.248 |
| SWHC × Field history | 0.033 | 0.857 | 0.074 | 0.786 | 5.083 | **0.025** | 1.527 | 0.218 |
| SWHC × Cultivar | 0.639 | 0.428 | 2.586 | 0.115 | 5.889 | **0.016** | 2.966 | 0.086 |
| Field history× Cultivar | 1.370 | 0.248 | 0.995 | 0.324 | 8.184 | **0.005** | 33.50 | **0.000** |
| SWHC × Irrigation | 1.021 | 0.318 | 2.419 | 0.127 | 4.334 | **0.038** | 0.877 | 0.350 |
| Field history × Irrigation | 1.571 | 0.217 | 0.433 | 0.514 | 0.304 | 0.582 | 22.30 | **0.000** |
| Cultivar × Watering | 0.051 | 0.822 | 0.010 | 0.921 | 9.011 | **0.003** | 11.39 | **0.001** |
| SWHC × Field history × Cultivar | 0.300 | 0.587 | 0.179 | 0.675 | 2.528 | 0.113 | 34.20 | **0.000** |
| SWHC × Field history × Irrigation | 0.090 | 0.766 | 0.127 | 0.723 | 3.272 | 0.072 | 47.18 | **0.000** |
| SWHC × Cultivar × Irrigation | 0.244 | 0.624 | 0.161 | 0.691 | 8.210 | **0.004** | 11.00 | **0.001** |
| Field history× Cultivar × Irrigation | 0.029 | 0.865 | 0.184 | 0.670 | 0.109 | 0.741 | 1.746 | 0.187 |
| SWHC × Field history × Cultivar × Irrigation | 0.029 | 0.865 | 0.074 | 0.787 | 2.497 | 0.115 | 1.132 | 0.288 |

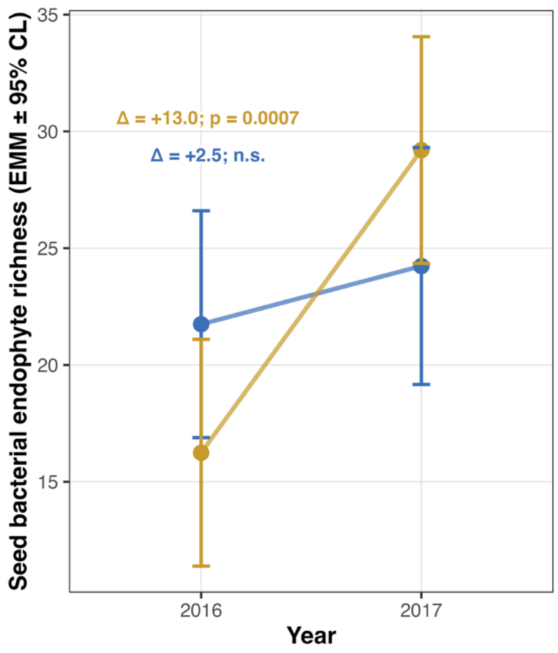

**Figure S1.** Estimated marginal means (EMMs ±95% CI) of seed endophyte richness in AC Nass wheat between dry (25–50 %) and wet (75–100 %) rainfall groups in 2016 and 2017. Blue color is devoted to the wet and gold color to the dry rainfall regimes. Error bars represent 95 % confidence limits derived from the fitted linear model.

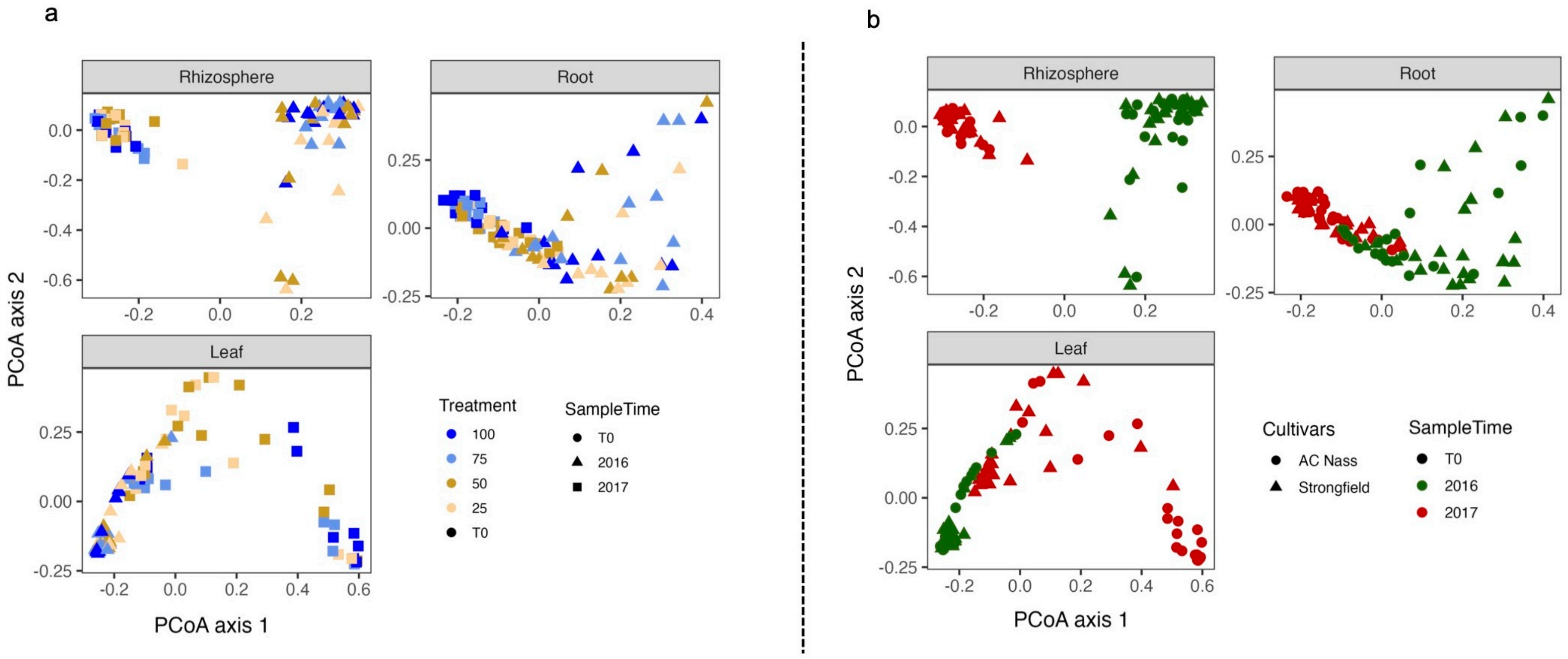

**Figure S2. Principal coordinate analysis (PCoA) of bacterial communities in compartments across rainfall treatments, cultivars, and years**. **a**, Community structure of bacterial assemblages in rhizosphere, root, and leaf samples of AC Nass, based on rainfall treatment (100, 75, 50, 25% of ambient) and sampling year (T0, 2016, 2017). **b**, Bacterial community structure in rhizosphere, root, and leaf samples grouped by cultivar (AC Nass, Strongfield) and sampling year (T0, 2016, 2017).

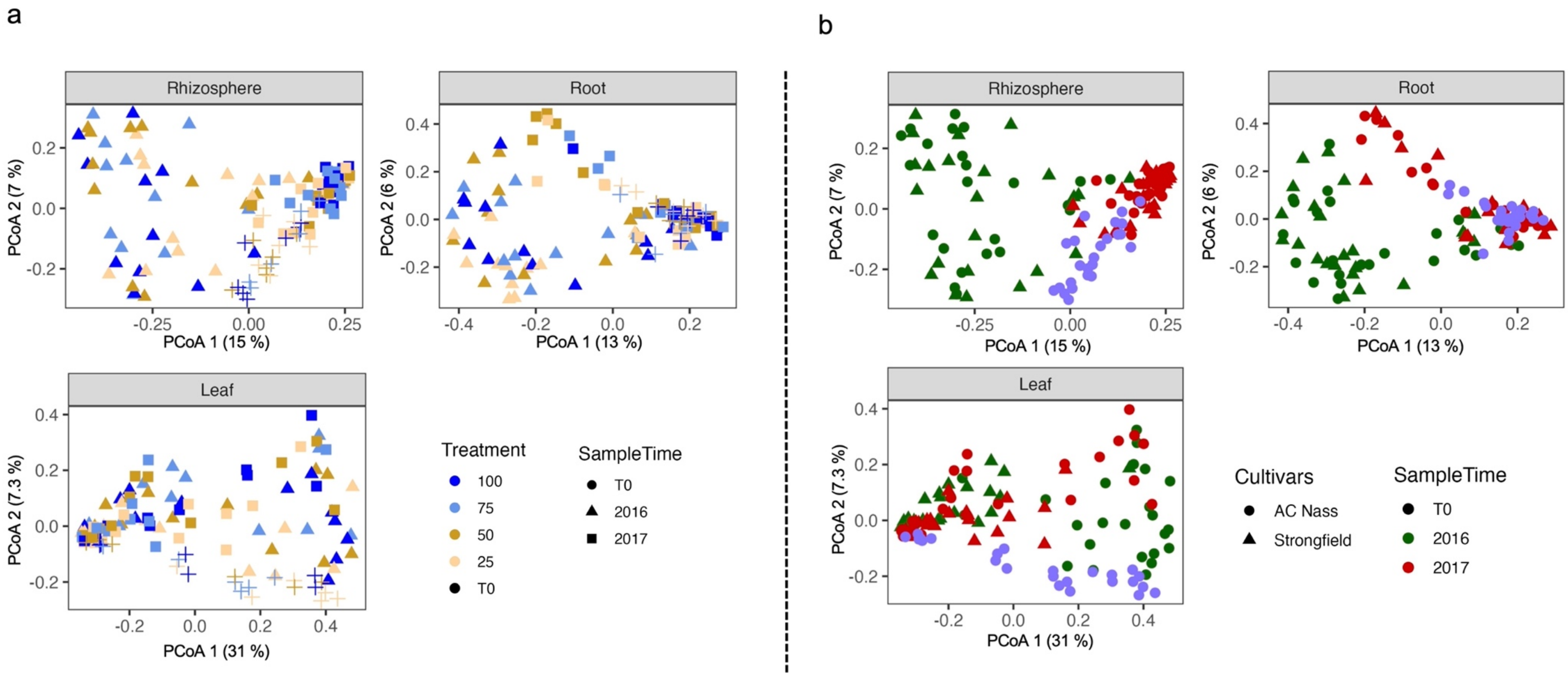

**Figure S3. Principal coordinate analysis (PCoA) of fungal communities in compartments across rainfall treatments, cultivars, and years**. **a**, Community structure of fungal assemblages in rhizosphere, root, and leaf samples of AC Nass, based on rainfall treatment (100, 75, 50, 25% of ambient) and sampling year (T0, 2016, 2017). **b**, Fungal community structure in rhizosphere, root, and leaf samples grouped by cultivar (AC Nass, Strongfield) and sampling year (T0, 2016, 2017).

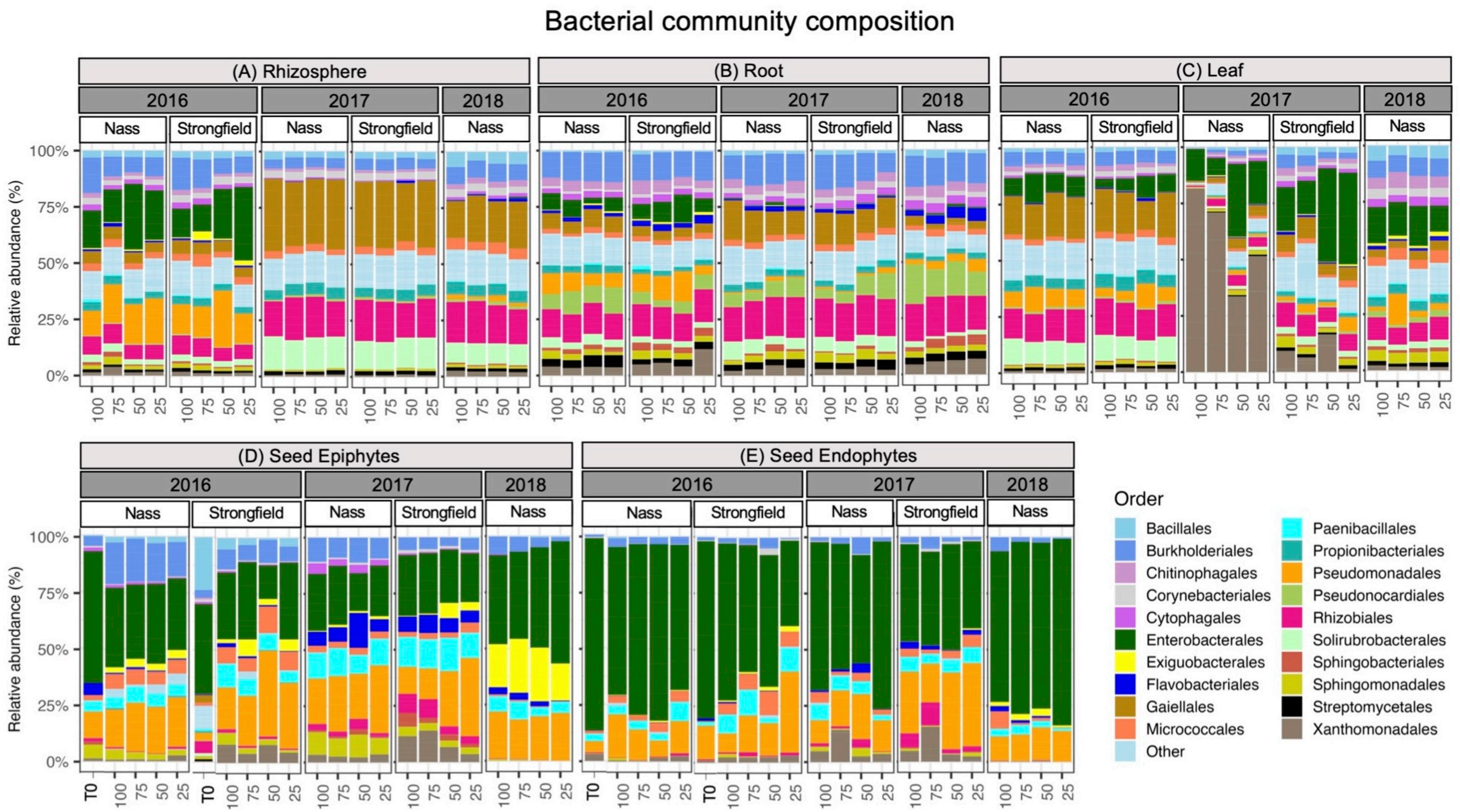

**Figure S4. Order-level bacterial community composition across plant compartments, cultivars, rainfall treatments, and years**. **a–c**, Relative abundances of bacterial orders in rhizosphere (a), root (b), and leaf (c) samples of AC Nass and Strongfield across rainfall treatments (100, 75, 50, 25% of ambient) and years (2016–2018). **d–e**, Relative abundances of bacterial orders in seed epiphytes (d) and endophytes (e).

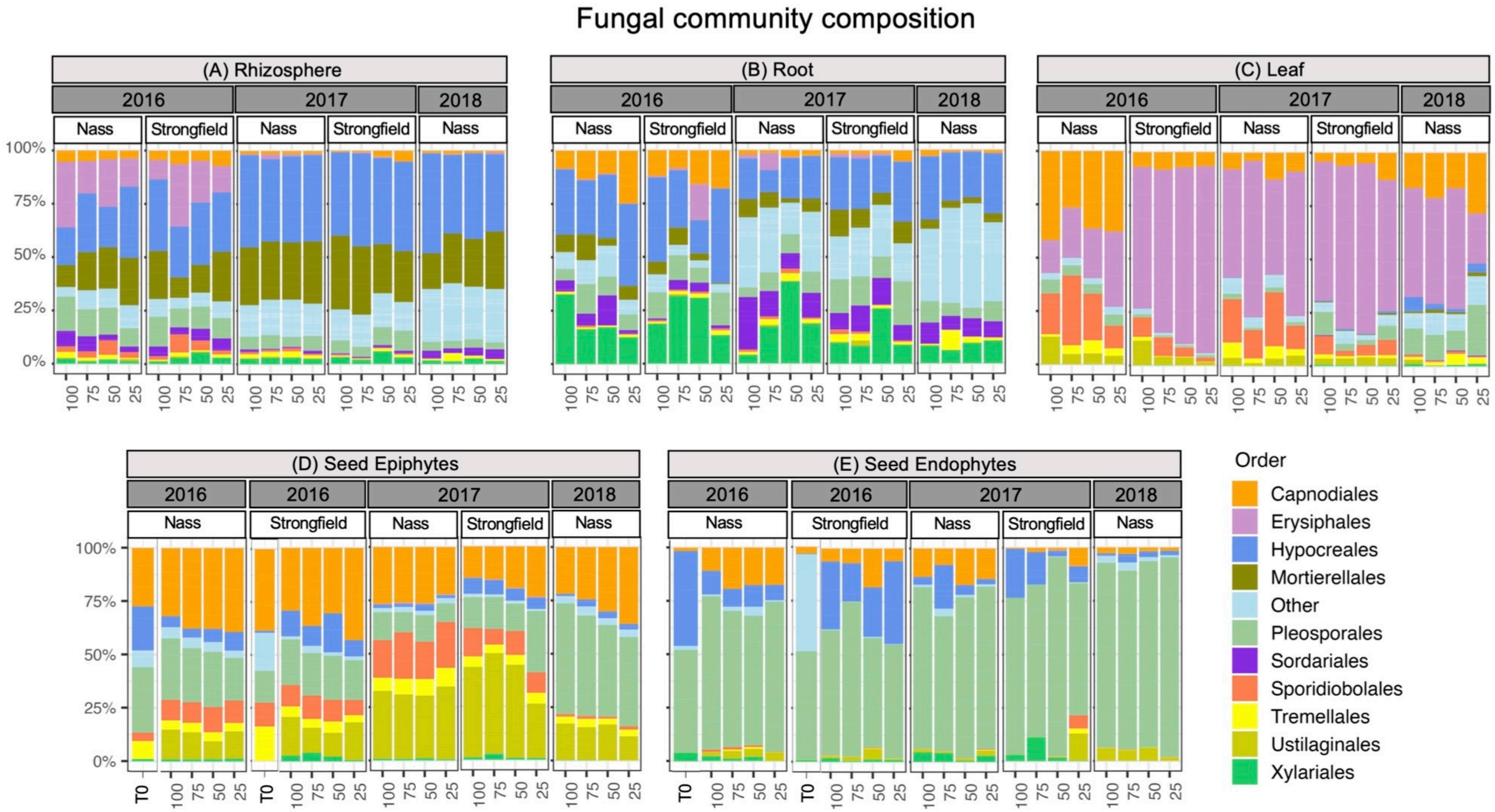

**Figure S5. Order-level fungal community composition across plant compartments, cultivars, rainfall treatments, and years**. **a–c**, Relative abundances of fungal orders in rhizosphere (a), root (b), and leaf (c) samples of AC Nass and Strongfield across rainfall treatments (100, 75, 50, 25% of ambient) and years (2016–2018). **d–e**, Relative abundances of fungal orders in seed epiphytes (d) and endophytes (e).

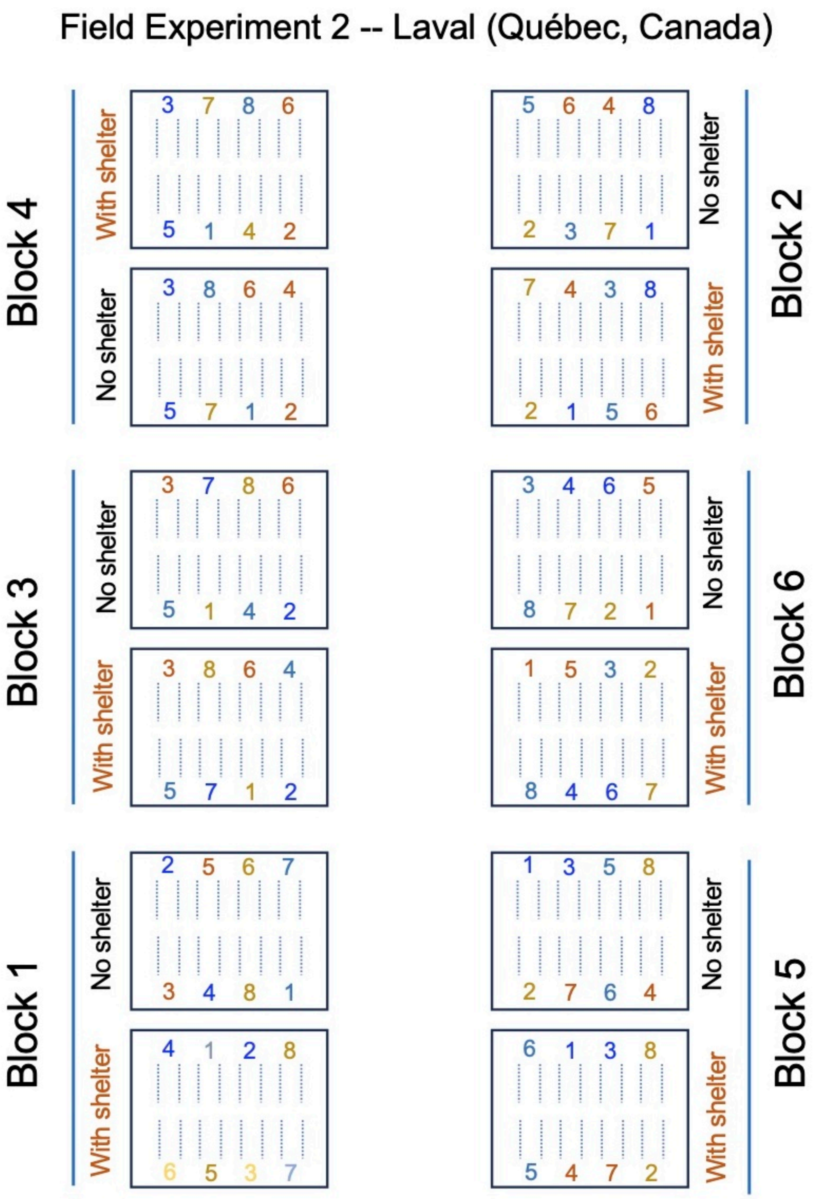

**Figure S6.** Experimental layout of the rain-out shelter trial (Experiment 2) across six blocks, with paired sheltered (reduced rainfall) and unsheltered (ambient rainfall) plots established for both cultivars (AC Nass, Strongfield) and for seeds originating from four precipitation histories in Experiment 1 (100, 75, 50, 25%).

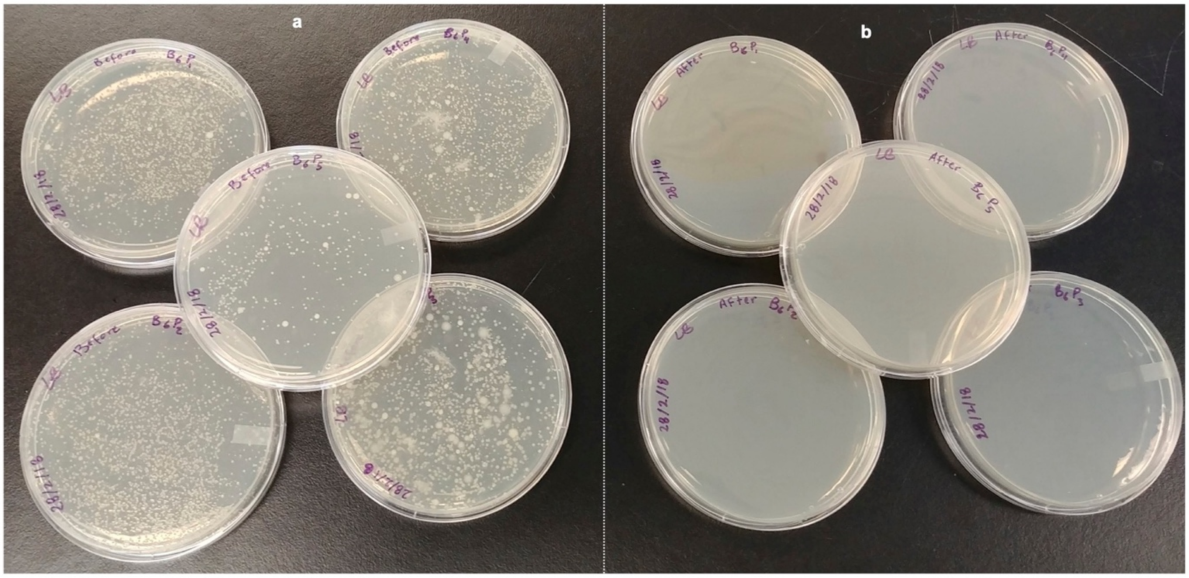

**Figure S7. Validation of surface sterilization protocol for seed microbiota extractions**. **a**, Bacterial growth on LB agar plates following imprinting of wheat seeds before surface sterilization, showing abundant microbial colonies derived from seed epiphytes. **b**, Absence of bacterial growth after surface sterilization, confirming effective removal of epiphytic microbes prior to extraction of seed endophytes.
